# Transcriptomic analysis of N-terminal mutated *Trypanosoma cruzi* UBP1 knockdown underlines the importance of this RNA-binding protein in parasite development

**DOI:** 10.1101/2023.12.07.570581

**Authors:** Karina B. Sabalette, Vanina A. Campo, José R. Sotelo-Silveira, Pablo Smircich, Javier G. De Gaudenzi

## Abstract

During its life cycle, the human pathogen Trypanosoma cruzi must quickly adapt to different environments, in which the variation in the gene expression of the regulatory U-rich RNA-binding protein 1 (TcUBP1) plays a crucial role. We have previously demonstrated that the overexpression of TcUBP1 in insect-dwelling epimastigotes orchestrates an RNA regulon to promote differentiation to infective forms. In an attempt to generate TcUBP1 knockout parasites by using CRISPR-Cas9 technology, in the present study, we obtained a variant transcript that encodes a protein with 95% overall identity and a modified N-terminal sequence. The expression of this mutant protein, named TcUBP1mut, was notably reduced compared to that of the endogenous form found in normal cells. TcUBP1mut-knockdown epimastigotes exhibited normal growth and differentiation into infective metacyclic trypomastigotes and were capable of infecting mammalian cells. We analyzed the RNA-Seq expression profiles of these parasites and identified 276 up- and 426 downregulated genes with respect to the wildtype control sample. RNA-Seq comparison across distinct developmental stages revealed that the transcriptomic profile of these TcUBP1mut-knockdown epimastigotes significantly differs not only from that of epimastigotes in the stationary phase but also from the gene expression landscape characteristic of infective forms. This is both contrary to and consistent with the results of our recent study involving TcUBP1-overexpressing cells. Together, our findings demonstrate that the genes exhibiting opposite changes under overexpression and knockdown conditions unveil key mRNA targets regulated by TcUBP1. These mostly encompass transcripts that encode for trypomastigote-specific surface glycoproteins and ribosomal proteins, supporting a role for TcUBP1 in determining the molecular characteristics of the infective stage.

## INTRODUCTION

The early-branching trypanosomes, such as *Trypanosoma cruzi* (the causative agent of Chagas disease) and *T. brucei* (the causative agent of African trypanosomiasis), are protozooan microorganisms that cause serious health problems in humans and domestic animals. In addition to their medical relevance, these pathogens are good models to examine novel mechanisms that control gene expression. Unlike most eukaryotes, these protists lack regulation at the level of transcription initiation for each gene. Contrarily, RNA polymerase II transcription is polycistronic, and transcript synthesis starts at a few locations on each chromosome [1–3]. During mRNA maturation, polycistronic pre-mRNA units are processed into monocistrons by two coupled reactions: 5′-*trans*-splicing and 3′-polyadenylation [4]. Because of these unusual genetic mechanisms, trypanosomatids primarily regulate protein levels through posttranscriptional processes. RNA-binding proteins (RBPs) that are bound to an mRNA determine its fate within the cell, affecting translation [5–7], nuclear export [8], mRNA stability [9], and mRNA processing [10]. RBPs can act as critical *trans*-acting factors at each of these regulatory points and mediate parasite differentiation in both *T. cruzi* and *T. brucei* [11–16]. The unicellular parasite *T. cruzi* has a complex life cycle characterized by alternating between replicative and infective forms in insect and mammalian hosts [17]. Intracellular amastigotes and bloodstream trypomastigotes are present in vertebrate hosts, whereas epimastigotes and metacyclic trypomastigotes are found in insect vectors of the family *Reduviidae* [18]. An important finding from the first RNA-Seq transcriptome and translatome of this parasite was that a key factor in the control of the gene expression profiles during *T. cruzi* differentiation is translation regulation [5]. The molecular mechanisms that could explain this regulation have been previously described by us and other authors [8, 19, 20].

In previous studies of our group, we also identified a superfamily of RNA-recognition motif (RRM)-type RBPs in the genome of kinetoplastid parasites [21]. One of the first reported trypanosome proteins with an RRM sequence is the trypanosome-exclusive *T. cruzi* U-rich RBP 1 (TcUBP1). This protein is a member of a subfamily composed of a small group of closely related proteins, including TcUBP2 [22], and plays a significant role in posttranscriptional gene regulation. The *TcUBP1* and *TcUBP2* genes have been found to be present in a single dicistronic unit, separated by an intercistronic region of ∼3 kbp [23], which is processed into mature monocistrons by trans-splicing/polyadenylation events. Both the structure and function of TcUBP1 have been the subject of extensive research due to their importance in the biology of *T. cruzi*. This small RBP has been found to interact with other proteins involved in RNA metabolism [24–26], suggesting its involvement in larger ribonucleoprotein complexes that participate in RNA processing, transport, and translation. TcUBP1 binds specifically to U-rich sequences within 3′-untranslated regions and has been involved in the coordinated gene regulation of multiple *T. cruzi* transcripts termed posttranscriptional regulons [22, 27–29]. Keene was the first to establish the revolutionary paradigm of RNA regulons almost two decades ago [30, 31], and described that cells can coregulate subsets of transcripts with a common physiological function by recognizing structural and/or sequence RNA elements. Although many studies have provided support for the concept of RNA regulons, the strongest evidence is found in kinetoplastid parasites [32].

Overexpression and CRISPR-Cas9 knockout models of RBPs have been valuable tools in trypanosome research [33–35]. Overexpression has helped to reveal specific RNA interactions and has provided insights into the functional consequences of increasing RBP levels. In a recent study, we successfully conducted TcUBP1 overexpression experiments following RNA-Seq and found important results regarding the regulatory functions of TcUBP1 in the parasite life cycle [16]. Our findings showed that TcUBP1 overexpression facilitates a switch toward the profile expression of infectious trypomastigotes by increasing the mRNA levels and translation rates of an RNA regulon for cell-surface trypomastigote glycoproteins (which play critical roles in host-parasite interactions and immune evasion) [36]. Consequently, TcUBP1 has been involved in the control of developmental transitions, including the differentiation of *T. cruzi* into infective forms. The influence of UBP1 knockdown on the parasite transcriptome is unknown. In this study, we aimed to investigate the extent of the regulatory effects of UBP1 by analyzing the gene expression profile that triggers its downregulation in epimastigote samples. Hence, the RNA-Seq data integration of UBP1 overexpression and downregulation allowed a comprehensive analysis, shedding light on the function of this important RRM-type RBP, which is crucial for differentiation to infectious stages.

## METHODS

### Plasmid construction, parasite cultures and transfection

Culture conditions for *T. cruzi* CL-Brener cloned stock [18], parasite transfections parameters, viability and differentiation experiments were according to Sabalette *et al.* [36].

### CRISPR-Cas9 experiments

The endonuclease cleavage site was designed using CHOPCHOP-CRISPR/gRNA (https://chopchop.cbu.uib.no), with *TcUBP1*_CDS (TcCLB.507093.220) as the input sequence. The hit with the higher score and no predicted off-targets was selected as the target site, complementary to pos. 13-36 of the *TcUBP1*_CDS. Target sequence: 5′-TTGCTGCTGCTGCAGCTGCTGTAGTTGGGCAGTCTGGCCGTACGGATCGTACTGTGA AACC< >AACGGATTTGGCTCAT-3′, < > endonuclease site. See primers used in **S1 Table**.

### In vitro infections

Infection experiments were according to Sabalette *et al.* [36]. Briefly, Vero cells were plated onto round coverslips 24 h before infection. Infections were performed for 4 h with 2×10^6^ cell-derived trypomastigotes per coverslip. After infection, the cells were washed twice in 1 x PBS and incubated in fresh medium for additional 48 h to allow amastigote replication.

### Antibodies and Western blotting

The antibodies used in this work were polyclonal rabbit antibody reacting with RNA-binding of TcUBP1 (anti-RRM) and polyclonal mouse antibody reacting with the N-terminal portion of TcUBP1 (anti-NH2-UBP1) [24]. Protein samples were resolved by SDS-PAGE gels 12.5%, transferred onto Hybond C nitrocellulose membrane (GE Healthcare), probed with primary antibodies anti-RRM diluted 1:300, and developed using horseradish peroxide-conjugated antibodies and Supersignal WestPico Chemiluminescent Substrate (Thermo Scientific). Protein expression values in KD epimastigote samples (n =3) were determined relative to WT controls by Western blotting analysis of TcUBP1 levels normalized to total protein loading, as measured by Coomassie Blue staining.

### Microscopy analysis

Immunofluorescence assays and epifluorescence experiments were according to Sabalette *et al.* [36]. The dilutions used for primary antibodies were 1:300 (anti-RRM) and 1:200 (anti-NH2-UBP1) and the dilution used for Alexa 568-conjugated goat anti-rabbit IgG (H+L) was 1:10,000 (Molecular Probes). Coverslips were mounted and photographed using a Nikon Y-FL fluorescence microscope.

### RNA preparation and RNA-Seq and overall quality of parameters of the transcriptomic data

RNA preparation, sequencing, and bioinformatic analysis were according to Sabalette *et al.* [16]. Three independent replicates of each condition were sequenced. A library was prepared at the BGI Americas Corporation. The samples were used for paired-end (PE) deep sequencing and the libraries were sequenced using 2 × 100 PE chemistry on DNBSeq platform to generate ∼3.9 GB of data per sample. After trimming of low-quality sequences, a total of ∼24M reads were obtained for each KD sample. The mapped read numbers obtained for UBP1mut-KD against the reference CL Brener Esmeraldo-like strain were 40,431,666 (replicate 1), 40,209, 302 (replicate 2) and 40,373,538 (replicate 3).

### Read processing and data analysis

Read processing and data analysis were performed as described [16].

### Functional annotation of gene lists

Gene ontology (GO) analysis was carried out for the DEGs from the TriTrypDB database (http://www.tritrypdb.org). All the genes for *T. cruzi* were taken as the reference set and the DEGs for both OE and KD were taken as the test set (consistent genes stabilized in OE and destabilized in KD and *vice versa*). The GO annotations were extracted and visualized as bubble charts using ggplot2 in R (www.r-project.org). Also, the results were visualized as word cloud using the Analyze Result tool from TriTrypDB.

## RESULTS

### Insertion of a stop codon in the *TcUBP1* gene generated a downregulated N-terminal mutant in epimastigote cells

To further investigate the regulatory role of TcUBP1, we intended to generate a population of CL Brener parasites knockout for this RRM protein by using the CRISPR-Cas9 technique. We initially tested the system by transfecting epimastigote cells with the plasmid pROCK-Cas9-GFP, which constitutively and ectopically expresses *Streptococcus pyrogenes* Cas9 (spCas9), the endonuclease used to generate the cut in the DNA molecule. We analyzed the correct expression and localization of the nuclease by using fluorescence microscopy (**Fig 1A**) and verified that the population presented a normal morphology, which allowed us to rule out the possibility of spCas9 toxicity on parasites in our working conditions. This population was *in vitro* transcribed with the RNA guide transcript sequence (see Methods). The strategy was optimized by incorporating the hammerhead and Hepatitis Delta Virus ribozymes to the 5′ and 3′ ends of the guide RNA respectively [37], increasing transfection efficiency by reducing the final length of the RNA molecule (100 nt) (**S1A Fig**). Finally, to be able to select the parasites with the interrupted *TcUBP1* gene and obtain a knockout epimastigote culture, we used a DNA donor molecule for the hygromycin resistance gene flanked at both ends by sequences complementary to the target gene (**S1B Fig**). Aliquots from transfected cultures with guiding RNA and donor DNA, as well as the respective spCas9-GFP and wildtype (WT) parasites, were analyzed by PCR. As expected, samples corresponding to the population with interrupted *TcUBP1* amplified a PCR fragment of >1500 bp by using specific primers for *TcUBP1*, corresponding with the sum of the size of the endogenous UBP1 coding sequence (CDS) (675 nt) and the hygromycin resistance CDS (1023 nt), whereas, in the controls, the amplicon size corresponded to the *TcUBP1* sequence (**S1C Fig**). When using hygromycin-specific primers, the presence of this gene was observed in the population transfected with the complete system (spCas9-gRNA-donor DNA), and no band was detected in the control populations (**S1C Fig**). Next, to corroborate the lack of expression of TcUBP1 in the parasites modified by CRISPR-Cas9, a Western blot analysis was performed using specific antibodies. Unexpectedly, the experiment revealed the presence of a protein band with a molecular weight slightly lower than that of TcUBP1, whose expression level was approximately 30% of the signal relative to the endogenous protein in WT parasites (0.27 ± 0.11 times) (**Fig 1B**).

**Fig 1.**
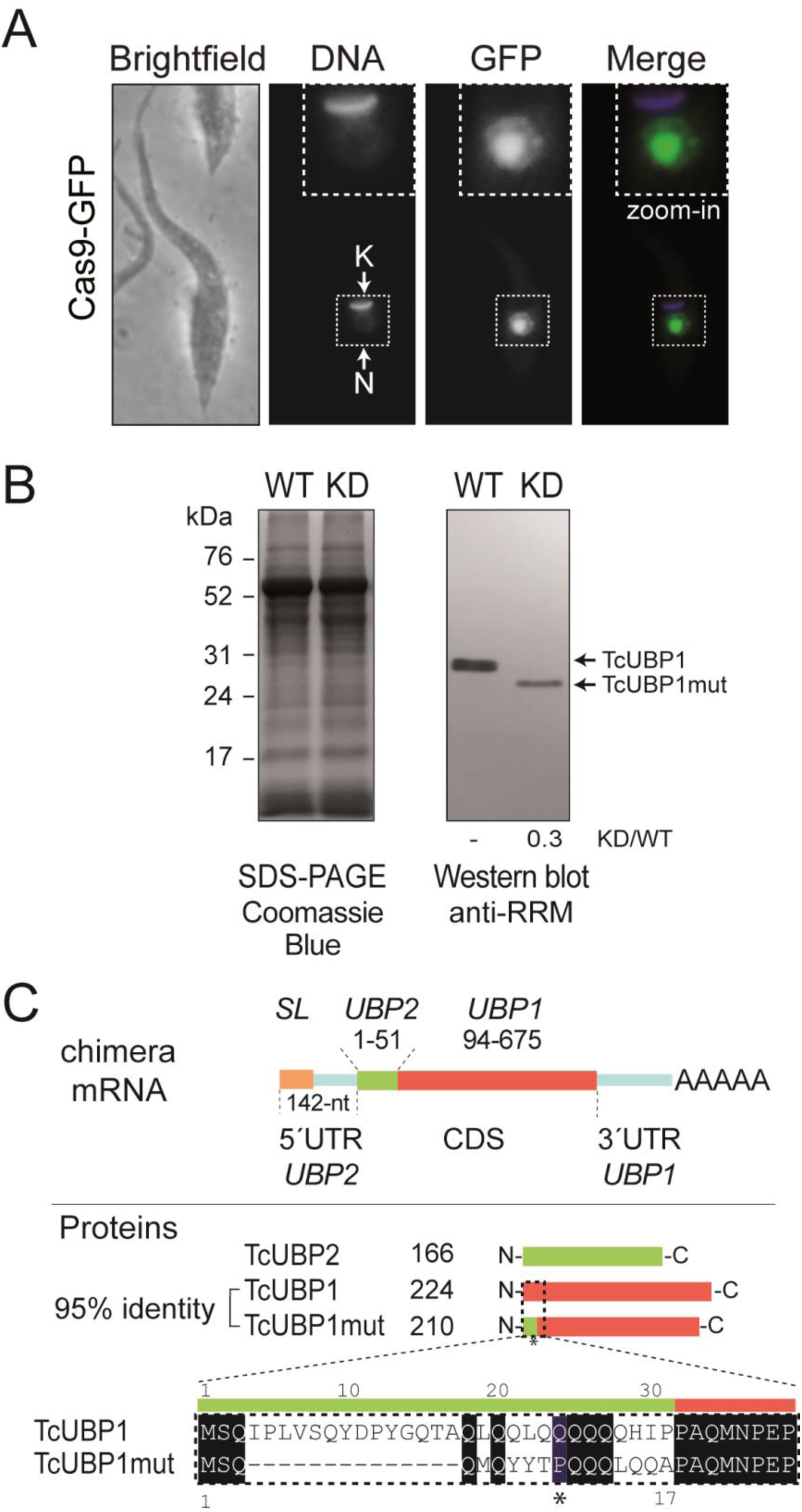
Diminished expression of the N-terminal mutant protein in TcUBP1mut knockdown parasites. *A*, *T. cruzi* controls parasites transfected with pROCK-Cas9-GFP. A representative image of a transfected epimastigote cell. Grayscale images of each channel are shown on the left, and a merged image showing expression of GFP (in *green*) and DAPI staining (in *blue*) for visualization of kinetoplast (K) and nuclear DNA (N) is shown on the right. *B*, Equal amounts of protein extracts from WT and KD epimastigotes were layered on each lane (see left panel, Coomassie staining gel). The blot was reacted with rabbit polyclonal anti-RRM antibodies. TcUBP1 and TcUBP1mut are indicated by arrows and relative quantification of the protein is indicated below each lane. *C*, Scheme of the chimera monocistronic RNA identified by sequencing in the UBP1mut population. Fragments of CDS for *TcUBP2* (green) and *TcUBP1* (tomato) are shown indicating the starting and ending positions. Scheme of TcUBP1, TcUBP2 and UBP1mut proteins and sequence alignment of the N-terminus of TcUBP1 and TcUBP1mut is shown. N, nucleus; K, kinetoplast DNA; *SL*, spliced leader sequence; AAAA, poly(A) tail.

We then attempted to find out whether the protein band revealed by Western blot corresponded to the *TcUBP1* CDS located downstream of the hygromycin STOP codon. To this end, an RT-PCR was performed using total RNA extracted from UBP1-CRISPR-Cas9-transfected parasites. cDNA was synthesized by using a T7(dT) primer, and subsequent PCRs were performed with different *TcUBP1* specific primers, combined with reverse T7 or forward *spliced-leader* (*SL*) oligonucleotides to detect a putative mutant *TcUBP1* monocistronic processed sequence (**S1D Fig**). Surprinsingly, DNA sequences from cDNA clones and RNA-Seq counts revealed the composition of a chimeric mRNA sequence coding for a hybrid TcUBP2-TcUBP1 product (**S1E** and **S2A Figs**). This sequence was possibly generated by an unusual rearrangement of the RNA molecule (see Discussion). The chimeric mRNA molecule codes for a protein with a 31-residue deletion at the N-terminal end compared to the TcUBP1 sequence and whose first 17 residues are the same as those of the TcUBP2 N-terminal end (**Fig 1C**).

Given that the nucleotides coded by *TcUBP2* are highly similar to the CDS of *TcUBP1*, when performing a pairwise alignment, the chimeric protein UBP2_(1-17)_/UBP1_Δ1-31_ showed 95.3% identity to the TcUBP1 sequence (ktuple 2, gap penalty 4, gap length 12) (**S2B Fig**). Considering this high sequence similarity, we refer to these modified parasites as mutant TcUBP1 knockdown (hereafter UBP1mut-KD) epimastigotes. Notice that another TcUBP1-ΔN mutant protein highly similar to UBP1mut (a TcUBP1_Δ1-34_ construct lacking the N-terminal glutamine-rich low-complexity sequence but containing the functional RRM and the C-terminal region of TcUBP1) has been previously characterized. *In vitro* studies have shown that this UBP1-ΔN protein can bind to RNA [24] and *in vivo* studies have shown that it can also be recruited to mRNA granules [38]. Therefore, the RNA binding activity and protein localization of TcUBP1 N-terminal deletion mutant expressed as GFP fusion protein have been shown to be identical to those of TcUBP1-GFP.

### Epimastigote parasites underexpressing an N-terminal mutant of TcUBP1 present normal morphology, viability, and differentiation to metacyclic trypomastigotes

We next analyzed the endogenously modified parasites with lower expression of the mutant TcUBP1, using microscopic techniques. This allowed us to observe the normal morphology of the epimastigotes, as well as the correct location of the nucleus and the kinetoplastid DNA (**Fig 2A**). In addition, we checked for the loss of TcUBP1 expression by incubating the KD samples with antibodies specific to TcUBP1 (anti-RRM and anti-NH2 TcUBP1) (**Fig 2A** and **B**). Figure 2 also shows the results of a normal WT sample (upper panels). As an experimental control, samples were incubated without primary antibodies, followed by incubation with secondary antibodies and detection reagents (**S3 Fig**).

**Fig 2.**
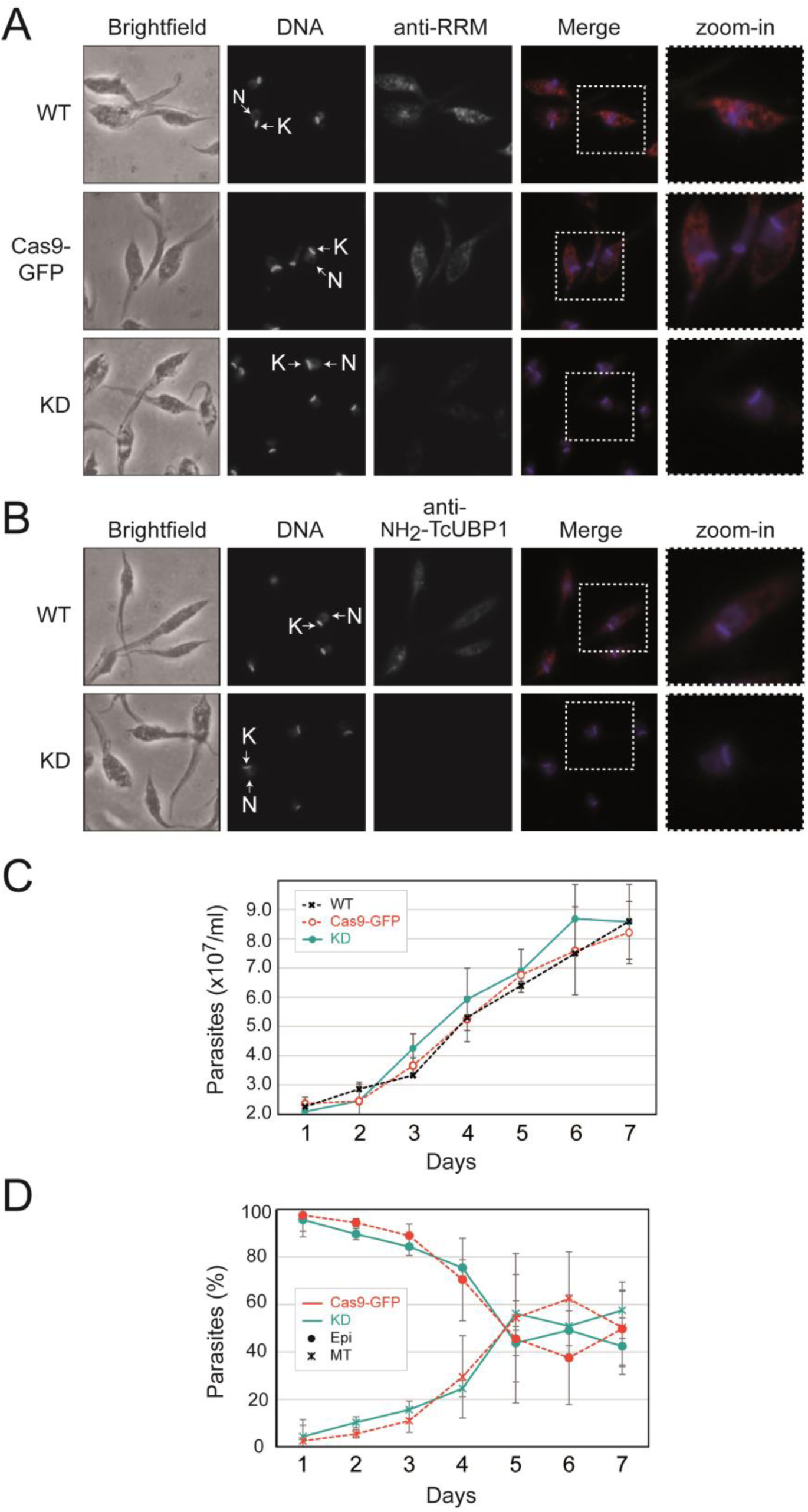
UBP1mut knockdown parasites have normal morphology and viability. *A*, Representative grayscale images of each channel are shown on the left for wildtype (WT), pROCK-spCas9-GFP (Cas9-GFP) and UBP1mut (KD) CRISPR-Cas9 transfected epimastigotes. Merged images showing expression of TcUBP1 (anti-RRM signal, in red) and DAPI staining (in blue) for visualization of kinetoplast (K) and nuclear DNA (N) are shown on the right. *B*, the key is as for *A*) using a mouse polyclonal N-terminal-specific antibody (anti-NH2-TcUBP1 signal in red) for visualization of TcUBP1 expression. *C*, Parasite growth curves of UBP1mut-KD (continuous line), wildtype populations (WT) and epimastigotes transfected with spCas9-GFP as experimental control (pointed lines, with markers in crosses and circles, respectively). Values of x10^7^ parasites/mL of culture medium are plotted according to the time at which a sample was taken (mean ± SD of three independent replicates). *D*, Differentiation from late-stationary phase cultured epimastigotes UBP1mut-KD (continuous lines) and those transfected with spCas9-GFP as controls (pointed lines). The percentage of the total population is plotted according to the time at which a sample was taken (mean ± SD of three independent replicates). Markers in circles refer to epimastigotes and crosses refer to metacyclic trypomastigotes.

We then evaluated axenic cultures of the UBP1mut-KD epimastigotes and observed that growth curves and differentiation to metacyclic trypomastigotes (when undergoing starvation stress) presented values similar to those of WT parasites and spCas9-GFP controls (**Fig 2C** and **D**). These results indicate that endogenous levels of TcUBP1 expression in replicative epimastigotes are not essential for parasite growth or differentiation to infective metacyclic trypomastigotes. We next performed Vero cell infection assays (**Fig 3A**) using CRISPR/Cas9 edited metacyclic trypomastigotes previously obtained. Results showed that the metacyclogenesis process (**Fig 2D**), the infection of mammalian cells (**Fig 3B**), and the subsequent differentiation and replication of intracellular amastigotes occurred normally (**Fig 3C**). The experiments with the spCas9-GFP control populations proceeded normally and the results indicated no significant differences in the infection percentage or the number of amastigotes per cell in comparison with the TcUBP1mut-KD population (Student’s *t* test, *p* value > 0.05).

**Fig 3.**
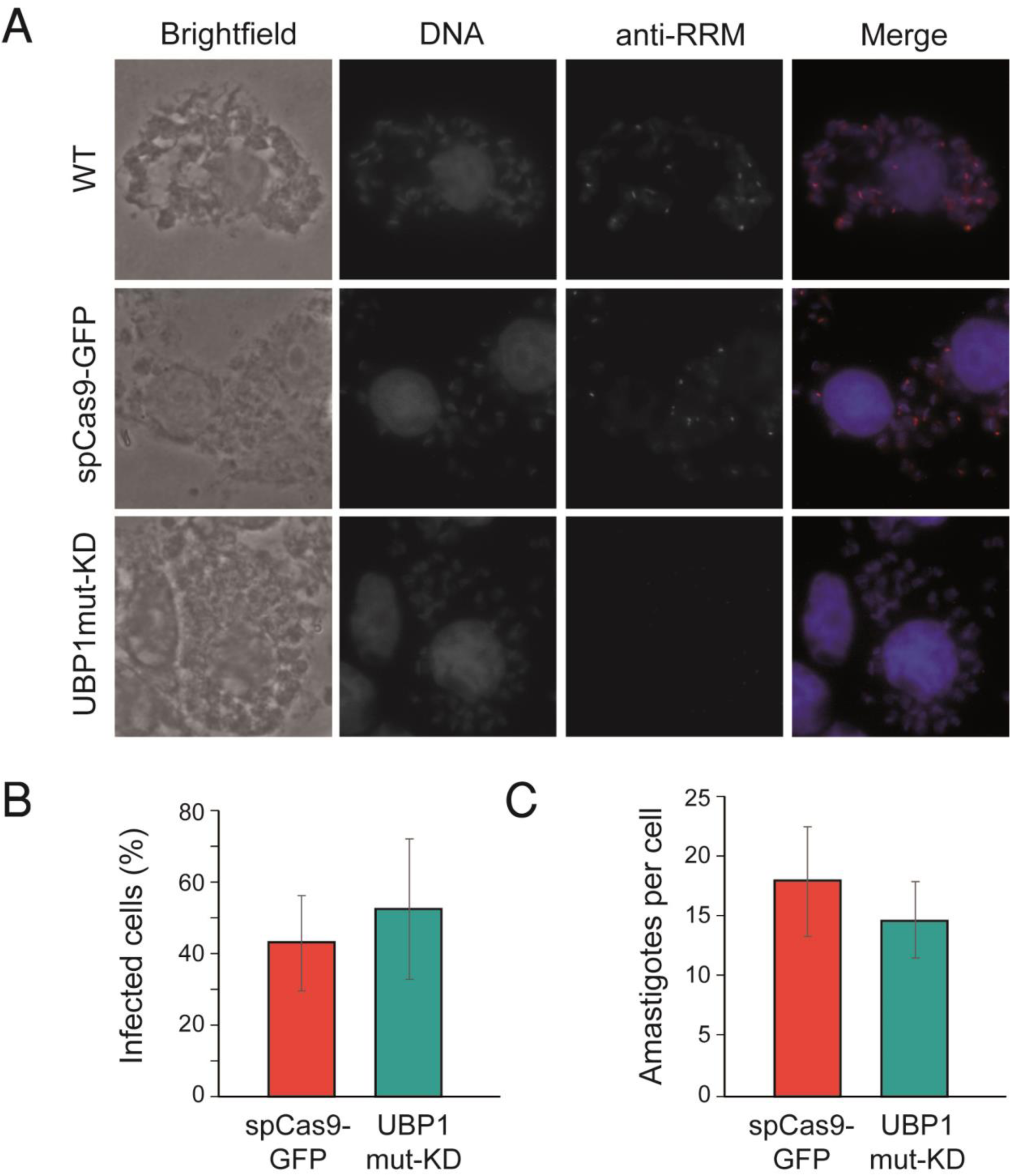
UBP1mut knockdown parasites lose the ability to differentiate from amastigotes to trypomastigotes. *A*, Representative photographs of infected cells with wildtype metacyclic trypomastigotes (WT), derived from epimastigotes transfected with spCas9-GFP or UBP1mut-KD. DAPI staining (in blue), TcUBP1 expression (anti-RRM signal in red). The images were taken four days after the infection. *B*, Quantification of number of infected cells obtained after 12 h from infections carried out with metacyclic trypomastigotes derived from spCas9-GFP or UBP1mut-KD samples (in percentage). *C*, Quantification of intracellular amastigotes obtained after 12 h from infections carried out in *B* (in amastigotes/cell). The values are expressed as the means of three independent experiments with the corresponding standard deviation bars.

### TcUBP1mut knockdown and full-length overexpression in epimastigotes have partly opposing effects on the parasite transcriptome

Our investigation continued to explore potential regulatory mechanisms that may be impacted by the downregulation of TcUBP1. Considering that this protein is involved in the stabilization or degradation of multiple mRNAs of different natures, we wondered whether the reduced TcUBP1 levels might play a role in gene expression. To achieve this objective, we conducted an RNA-Seq experiment from samples overexpressing the full-length TcUBP1 (OE), a mutated UBP1-knockdown (KD), and wildtype (WT) epimastigotes. We first verified the correct relationship between replicas and variability between experiments (**Fig 4A**). The expression profiles of these OE samples have been recently examined in our lab [16]. Now, we expanded the transcriptome analysis to the KD samples. As expected, TcUBP1 was >6 log2 higher in the OE samples (false discovery rate (FDR)-adjusted *p* value = 2.09E-84) and ∼-1 log2 lower in the KD samples (FDR-adjusted *p* value = 0.021), with respect to WT control parasites. Thus, the UBP1 downregulated expression value in KD parasites was comparable to the protein levels of UBP1mut detected in the Western blots (**Fig 1B**, lane KD: ∼0.3X or three-fold downregulated).

**Fig 4.**
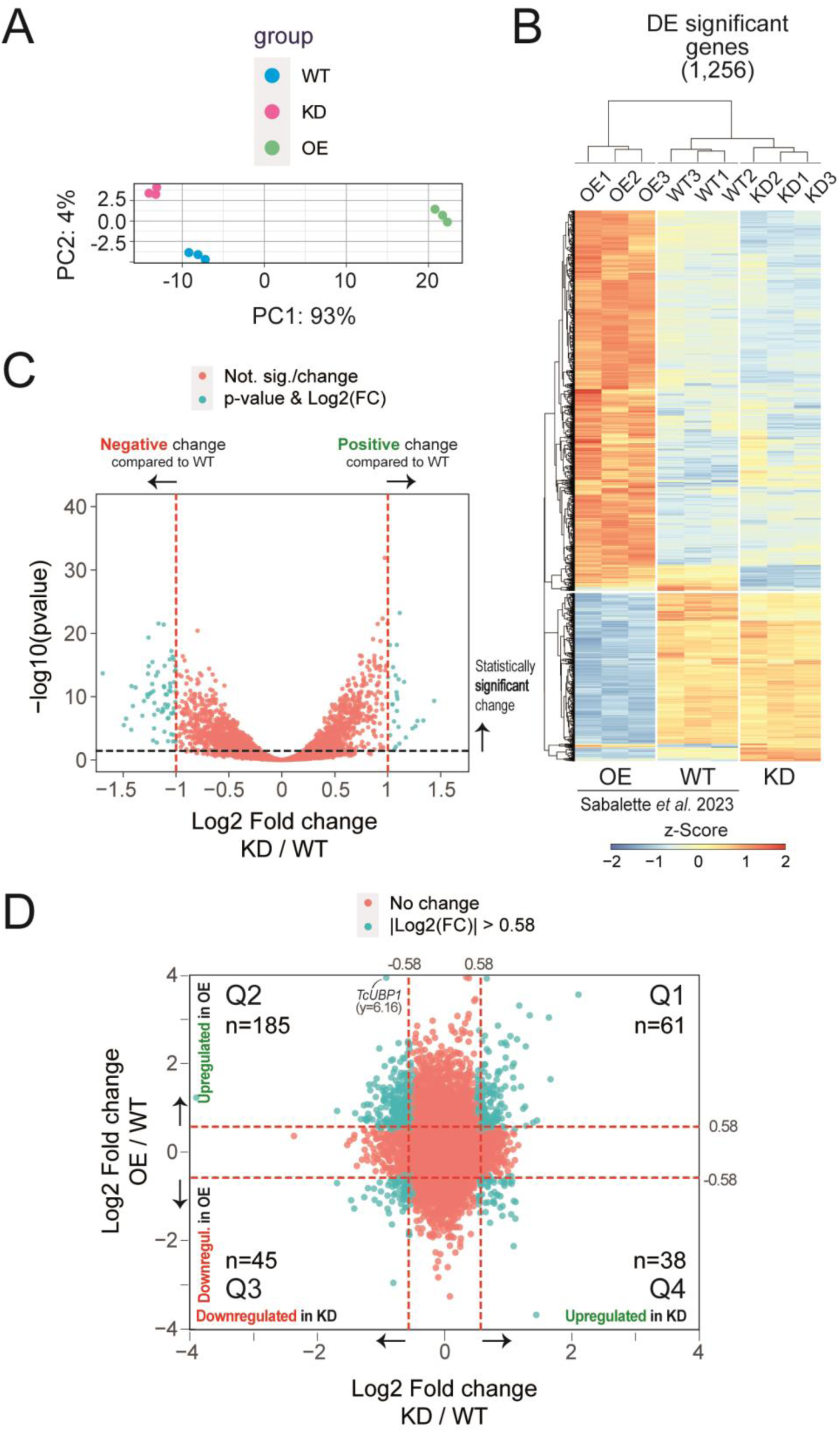
Hierarchical clustering of genes in OE, KD and WT samples defined by DESeq2. ***A***, PCA plot displaying all 9 samples along PC1 and PC2, which describe 93% and 4% of the variability, respectively, within the expression data set. PC analysis was applied to normalized (reads per kb of transcript per million mapped reads) and log-transformed count data. ***B***, Heatmap and complete linkage clustering using all replicates per group, 1,230 significant genes with |log2 fold change | > 1 were clustered. Groups on the vertical represent clustered genes based on gene expression with an FDR-adjusted *p* value lower than 0.05, the horizontal line represents a single gene and the color of the line indicates the gene expression. The Z-score scale bar represents relative expression +/- SD from the mean. ***C***, volcano plot showing the differential expression analysis of genes in UBP1mut-KD and WT parasites. Tomato and cyan dots show nonsignificant and significant DEGs, respectively. ***D***, Four-quadrant scatter plot showing the log2 fold change values in UBP1-OE (y axis) *versus* UBP1mut-KD (x axis). Quadrants Q2 and Q4 exhibit consistent gene profiles, as discussed in the text. Tomato and cyan dots show nonsignificant and significant DEGs, respectively.

After validating the expression of TcUBP1 in the KD dataset, we started the analysis of the expression profiles. We obtained a total of 6,487 significant genes with FDR-adjusted *p* value < 0.05 and 1,256 differentially expressed genes (DEGs) with |log2 fold change | > 1. At a large-scale level, we observed a higher number of genes whose abundance was lower than that of controls as a result of TcUBP1 knockdown. The expression patterns for the DEGs in the control, OE and KD samples are shown in **Fig 4B**. The light blue, white, and orange colors indicate less expressed, medium-level and highly expressed genes, respectively (**Fig 4B**). In particular, 26 genes were two-fold upregulated and 67 genes were two-fold downregulated in TcUBP1mut-KD samples relative to the WT (|log2 fold change| > 1, FDR-adjusted *p* value < 0.05; **S1 File**). A volcano plot of gene expression in UBP1mut-KD *versus* WT parasites is shown in **Figure 4C**, where significantly expressed genes are separated from non-significantly expressed genes by different color codes. By expanding the analysis of the expression profile of regulated genes using a criterion of at least 1.5-fold (|log2 fold change| > 0.58, FDR-adjusted *p* value < 0.05), we confirmed that the overall effect of UBP1mut-KD is primarily to destabilize the parasite transcriptome. We observed that 276 mRNAs increased in abundance, while 426 mRNAs decreased (**S2 File**). This result is consistent with the antagonistic effect observed after TcUBP1 overexpression [16]. Also, the Top 20 list with the most differentially over- or underexpressed genes from the OE, KD and WT epimastigotes (based on fold changes values) is shown in **S4 Figure**. Expression analysis using a four-quadrant scatter plot of fold change values between OE *versus* KD revealed their partly opposite expression patterns, evident in quadrants 2 and 4 (**Fig 4D**). Specifically, in quadrant 2 (Q2) 185 genes met the criteria of log2 fold change OE/WT > 0.58 & log2 fold change KD/WT < −0.58 and FDR-adjusted *p* value < 0.05, while cuadrant 4 (Q4) comprised 38 genes meeting the criteria log2 fold change OE/WT < −0.58 & log2 fold change KD/WT > 0.58 (**S3 File**). In light of this, we will refer to these genes as ‘robust genes’ or ‘consistent genes’ (see below).

We next evaluated how the variations of the TcUBP1 protein abundance levels in OE and KD samples are correlated with both the number of affected genes and the levels of variation they show. We found that several robust genes are consistently regulated, either increasing or decreasing their mRNA levels depending on the abundance of TcUBP1 and being oppositely affected in OE and KD conditions (**Table 1** and intersected genes in the Venn diagrams of **Fig 5A**). By analyzing the nature of these robust gene groups that were consistently affected, we identified 10 genes that were stabilized by TcUBP1, using a criterion of at least two-fold change (|log2 fold change| >1) compared with WT parasites (**Fig 5A**, right). As expected, four of these transcripts were mucins: TcCLB.509631.90 (MUCII_1), TcCLB.510275.330 (MUCII_2), TcCLB.504039.120 (MUCII_3), and TcCLB.508097.81 (MUCII_4), five were mucin-associated surface proteins: TcCLB.506677.70 (MASP_1), TcCLB.506763.260 (MASP_2), TcCLB.510377.134 (MASP_3), TcCLB.506501.350 (MASP_4), and TcCLB.507957.150 (MASP_5), and one was a putative peptidase-domain containing protein TcCLB.506755.50 (PPPDE) (**Table 1** and **Fig 5B**, right). On the other hand, six genes were found destabilized by TcUBP1 (**Fig 5A**, left), all coding for hypothetical proteins: TcCLB.503599.60 (HYPO_1), TcCLB.503599.70 (HYPO_2), TcCLB.503453.10 (HYPO_3), TcCLB.508125.40 (HYPO_4), TcCLB.511469.105 (HYPO_5), and TcCLB.5034534.4 (HYPO_6) (**Table 1** and **Fig 5B**, left).

**Fig 5.**
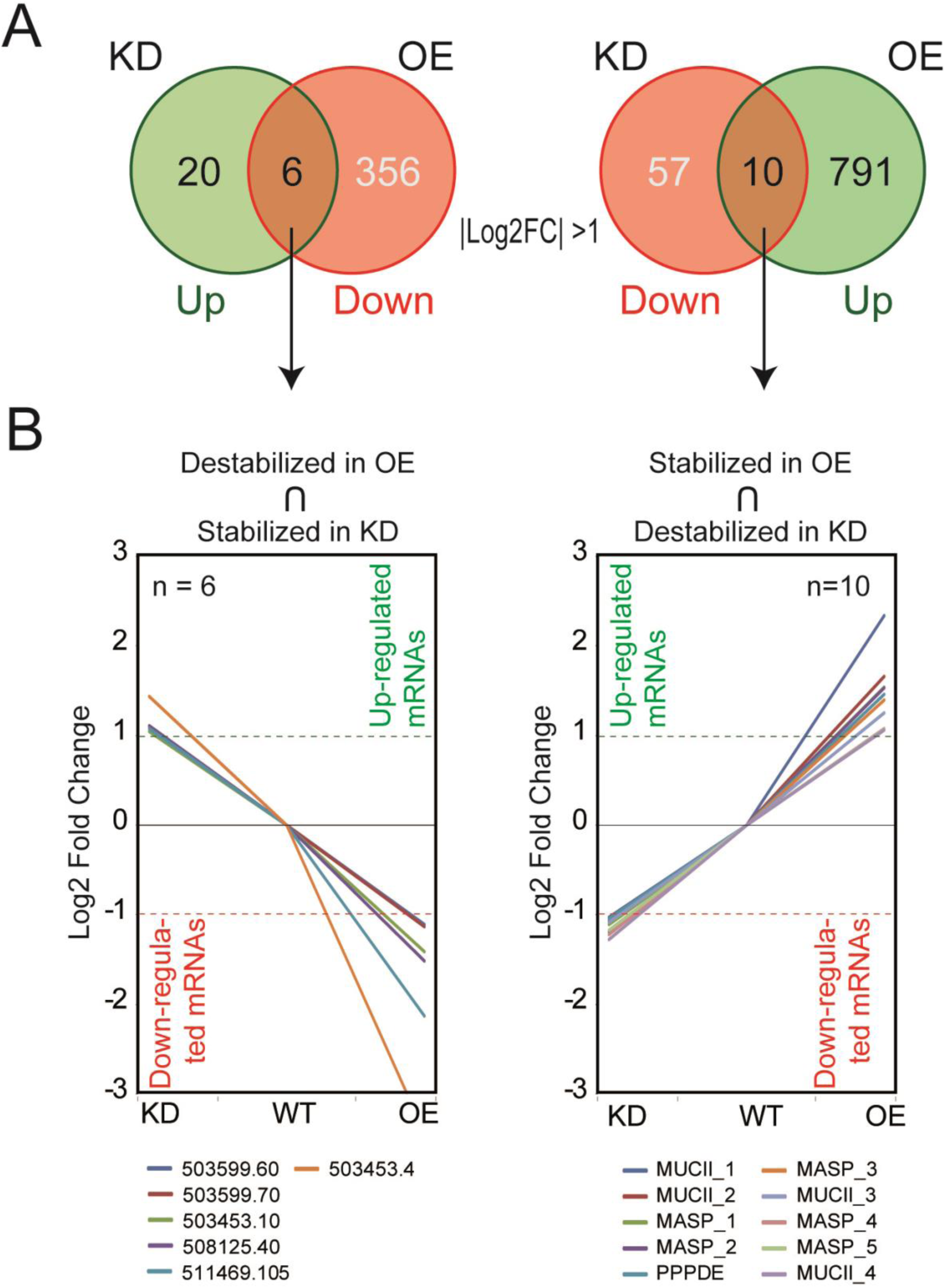
Transcripts whose abundance directly responds to TcUBP1 expression levels. ***A***, Venn diagrams showing the number of genes two times affected in each condition (OE and KD) with respect to the WT control (|log2 fold change| >1). In green, upregulated genes; in light red, downregulated genes; in orange, robust genes regulated in a coordinated way (mRNA abundance increased in one condition and decreased in the other). ***B***, Left, spaghetti plots showing fold enrichment of expression relative to the control sample of those genes whose abundance increases in epimastigotes KD and decreases in parasites overexpressing UBP1 (OE). Right, *vice versa*. The complete gene IDs and names of the genes are shown in **Table 1**. The pointed line marks a difference of ± 2X.

**Table 1.**
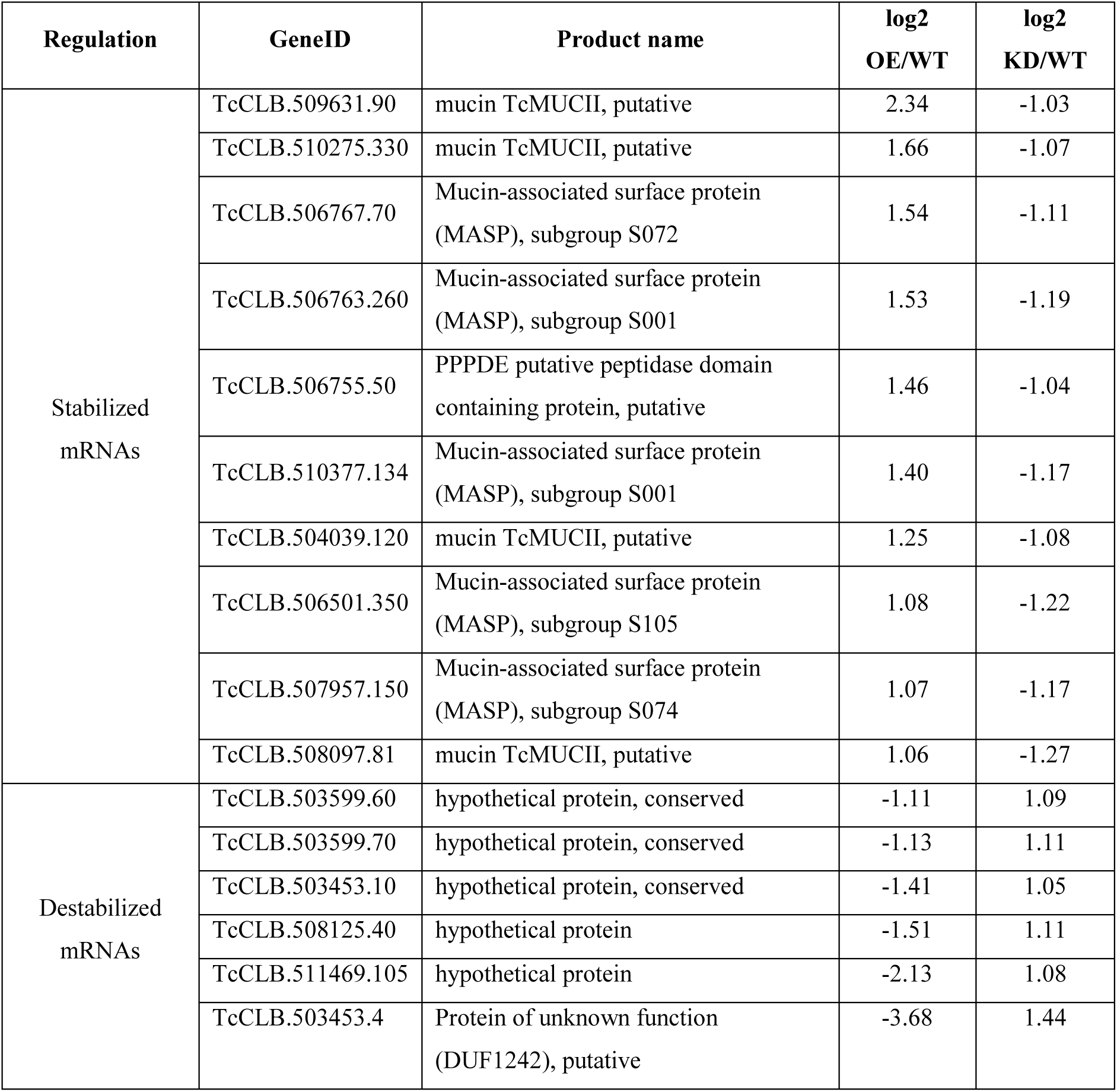
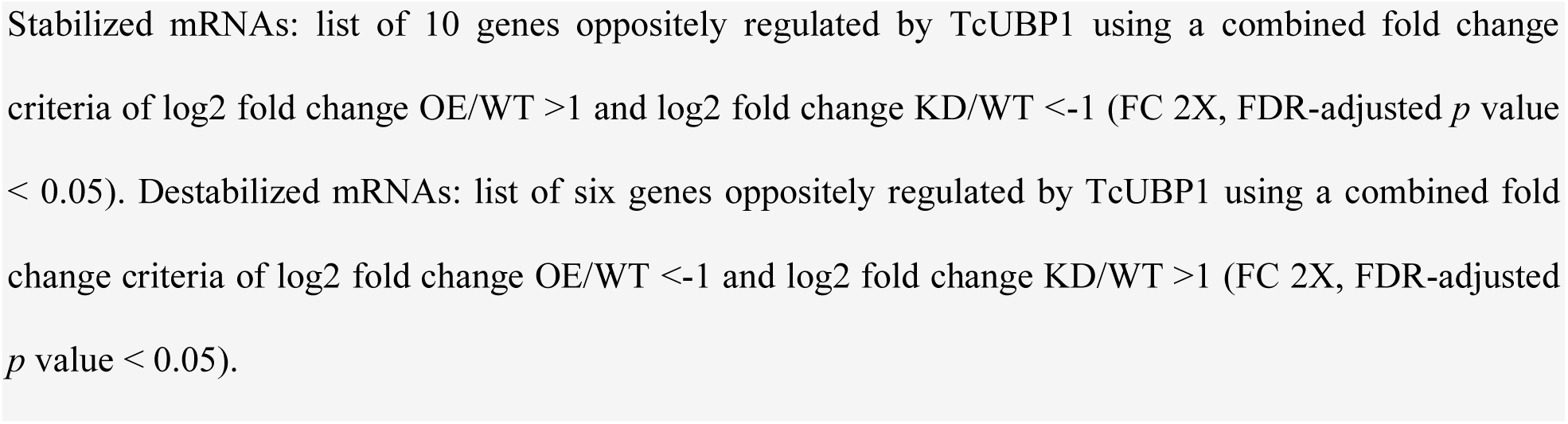
Oppositely affected genes after UBP1 overexpression and knockdown.

Revisiting the analysis depicted in Figure 4*D*, by using a less stringent expression cutoff of 1.5-fold change, we were able to identify 185 consistent genes up- and 38 downregulated by TcUBP1 (list A and B in **S3 File**, respectively and **S5 Fig**). Gene ontology (GO) analyses using TriTrypDB performed on these two lists, for the molecular function domain showed a distribution of 13 and 7 GO overrepresented terms, respectively (see the complete term distribution in **Table 2**). A plot for all the three GO domains, biological process, cellular component, and molecular function is presented in Figure 6 and revealed that genes upregulated in OE cells and downregulated in KD cells are involved in critical *T. cruzi* functions, such as alpha-sialidase activity (**Fig 6A**), and that genes upregulated in KD and downregulated in OE samples are mostly involved in translation and rRNA binding (**Fig 6B**). This analysis was also visualized using an enrichment word cloud for the four quadrants of Figure 4*D*. Upon reviewing the word cloud for Q1, no distinctive terms stand out prominently and in the word cloud for Q3, the term "calcium ion binding" barely emerges from the rest (**S6 Fig**), suggesting that quadrants Q2 and Q4 are most prominently enriched in a specific biological pathway.

**Fig 6.**
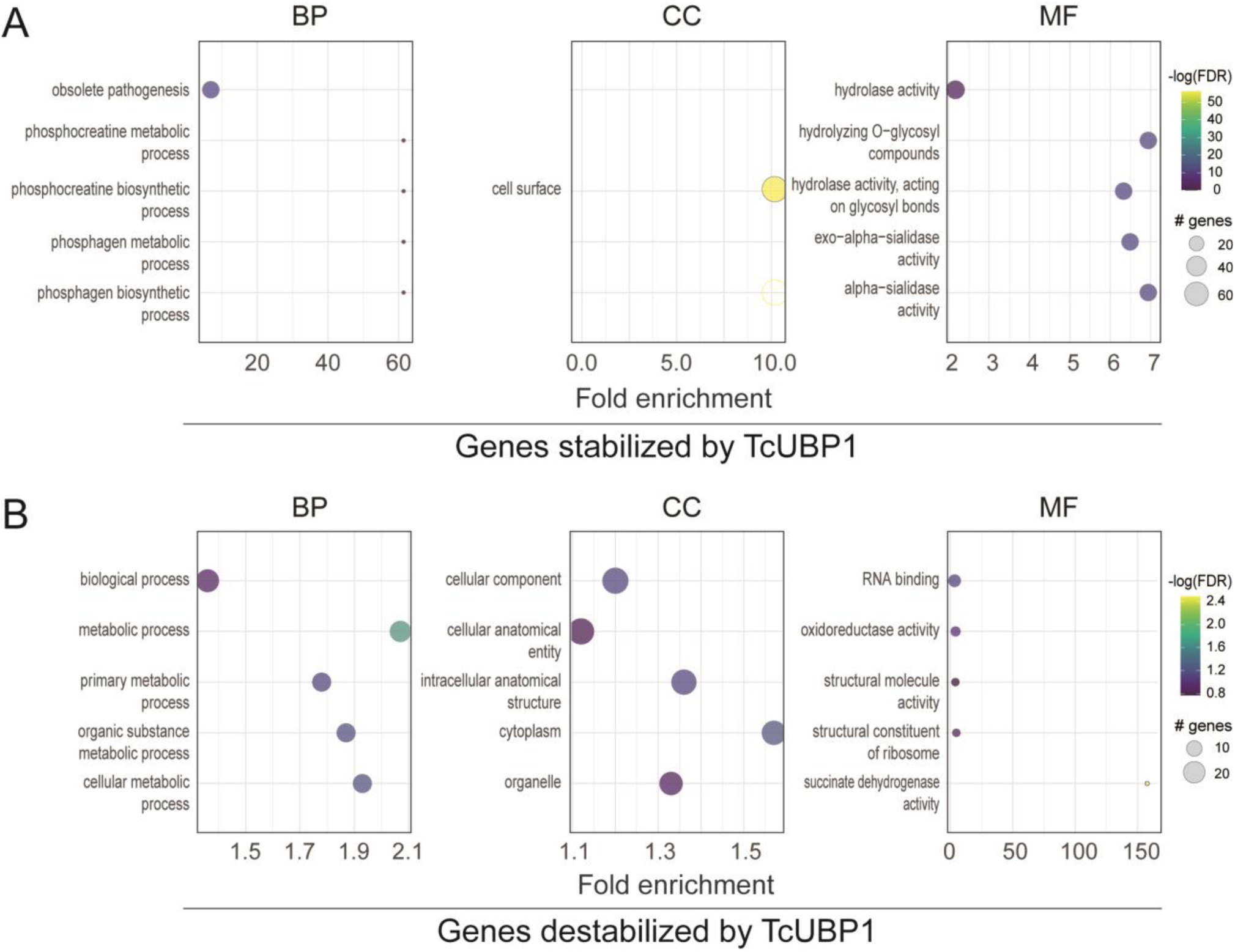
GO enrichment analysis of genes oppositely regulated in UBP1-OE and KD parasites. GO classification of DEGs using a criterion of at least 1.5-fold change (|log2 fold change| > 0.58); the graphs show up to 5 GO terms with most genes annotated and a word cloud of terms (only for GO domain: molecular function). BP, CC and MF charts indicated GO terms clustered in the biological process, cellular component and molecular function terms, respectively. The size of the dots diameter indicates the number of genes; color depth indicates significance; abscissa indicates enrichmen; and the ordinate indicates different pathways. *A*, 185 genes putatively augmented by TcUBP1: upregulated in OE and downregulated in KD samples. *B*, 38 genes putatively diminished by TcUBP1: upregulated in KD and downregulated in OE samples.

**Table 2.**
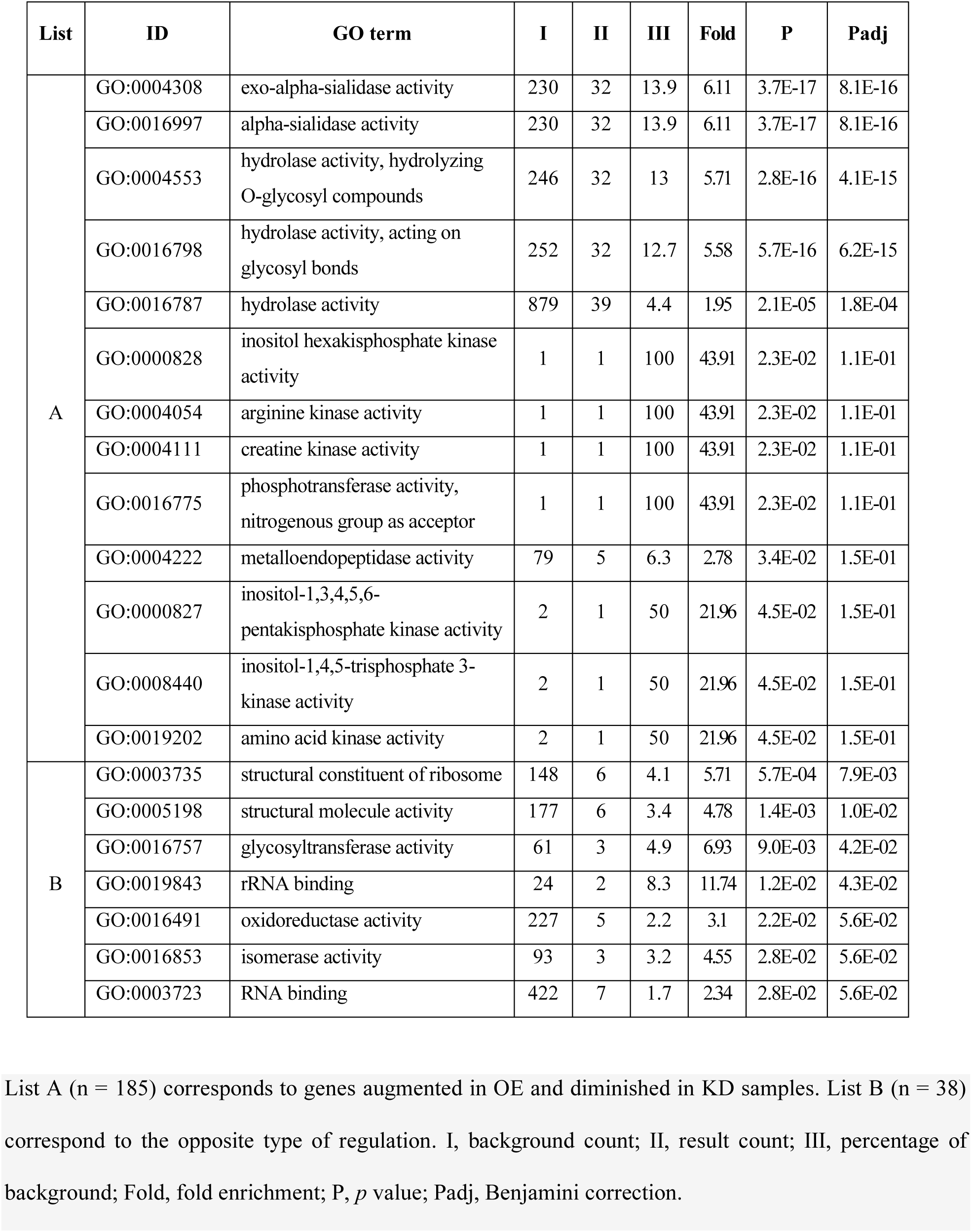
List of GO terms of the molecular function category enriched in the gene lists A and B.

### Clusters of transcripts coding for cell-surface trypomastigote glycoproteins and ribosomal proteins are directly influenced by TcUBP1 expression levels

By performing a comparative analysis of several functional gene groups, we further analyzed the expression profile triggered by TcUBP1. Based on the data presented in the section above, we manually classified 1,758 sequences obtained from UBP1-OE and KD parasites into 11 different categories: mucin-associated proteins (MASP), mucins (MUCI/II), the trans-sialidase/trans-sialidase like (TcS) superfamily, ribosomal proteins, protein phosphatases, protein kinases, heat-shock proteins, proteins involved in synthetic pathways, polymerases, RNA transcription, and transporter proteins. We then analyzed the abundance distribution of the transcripts in each category in both conditions (OE and KD) relative to the control (WT) by using violin plots showing expression values (log2 fold change OE/WT compared to KD/WT) (**Fig 7**). Results confirmed that the only group that showed significant enrichment in the KD samples with respect to the OE corresponded to ribosomal proteins, which are mostly expressed in replicative stages, whereas the groups of genes significantly enriched in the OE samples compared to KD are specific to the infective stages: *MASP*, *MUCI/II* and the *TcS* superfamily (Student’s *t*-test, \*\*\**p* value < 0.001). Moreover, proteins involved in synthetic pathways, protein kinases and phosphatases revealed significant differences between conditions (Student’s *t*-test, \**p* value < 0.05; \*\**p* value < 0.01). Finally, the categories of polymerase enzymes, heat-shock proteins, transporter proteins and transcription-associated proteins showed no significant differences as a consequence of TcUBP1 expression levels. Taken together, transcriptome analysis of both KD and OE parasites reveals that the expression of four mRNA families key for establishing a replicative or infective profile (*ribosomal proteins*, *MASP*, *MUCI/II* and *TcS*), respond directly and proportionally to TcUBP1 protein levels, indicating that this RBP plays a pivotal role in either triggering or not the differentiation process.

**Fig 7.**
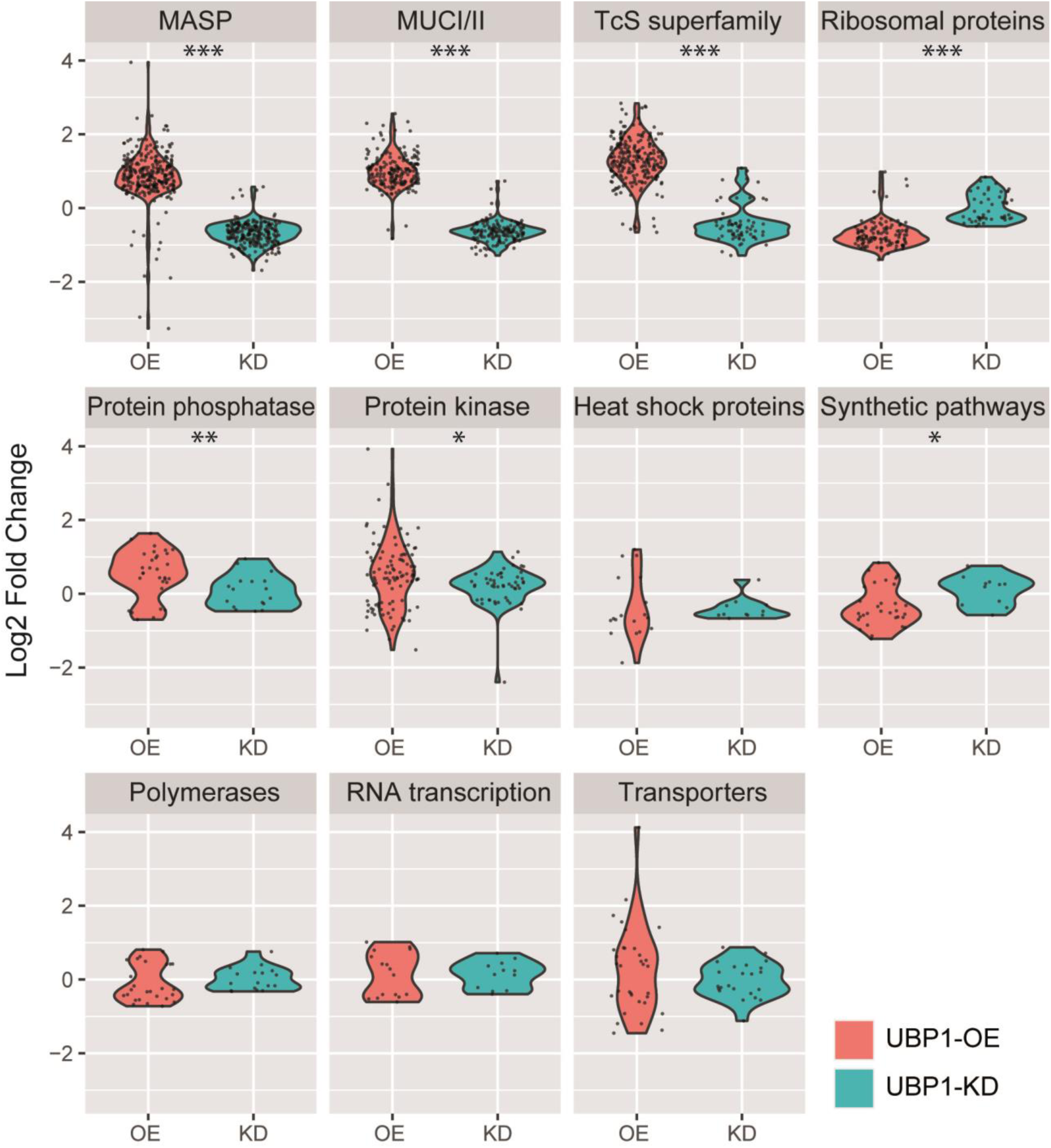
Violin plots displaying the expression distribution of the genes within 11 different functional categories in the OE or KD transcriptomes relative to the WT control (log2 fold change, FDR-adjusted *p* value < 10%). Categories in the figure are indicated at the top of each panel. Student’s *t*-test, * *p* value < 0.05,** *p* value < 0.01,*** *p* value < 0.001.

The most abundant cluster among the upregulated genes in the OE transcriptome was that coding for trypomastigote cell-surface glycoproteins [16]. Within the *TcS* superfamily group, we identified 42 significant robust genes (**Table 3**), whose abundance in both OE and KD conditions varied in a coordinated way, by either increasing or decreasing their mRNA levels depending on the abundance of TcUBP1 (|log2 fold change| >0.45, FDR-adjusted *p* value < 10%, **Fig 8A**). From this list, 29 genes (69%) are increased more than 1.5-fold in the OE experiment, and decreased in the KD condition (|log2 Fold change| >0.58, FDR-adjusted *p* value < 10%), a value that is restricted to nine genes, when considering those that decrease more than 1.75 times in relation to the WT control sample (|log2 Fold change| >0.8, FDR-adjusted *p* value < 10%; marked with black dots in **Fig 8A**). These results suggest that the mRNA abundances of members of the *TcS* superfamily are directly influenced by TcUBP1 levels, both increasing their expression when UBP1 protein levels are high and decreasing their expression when UBP1 protein levels are low.

**Fig 8.**
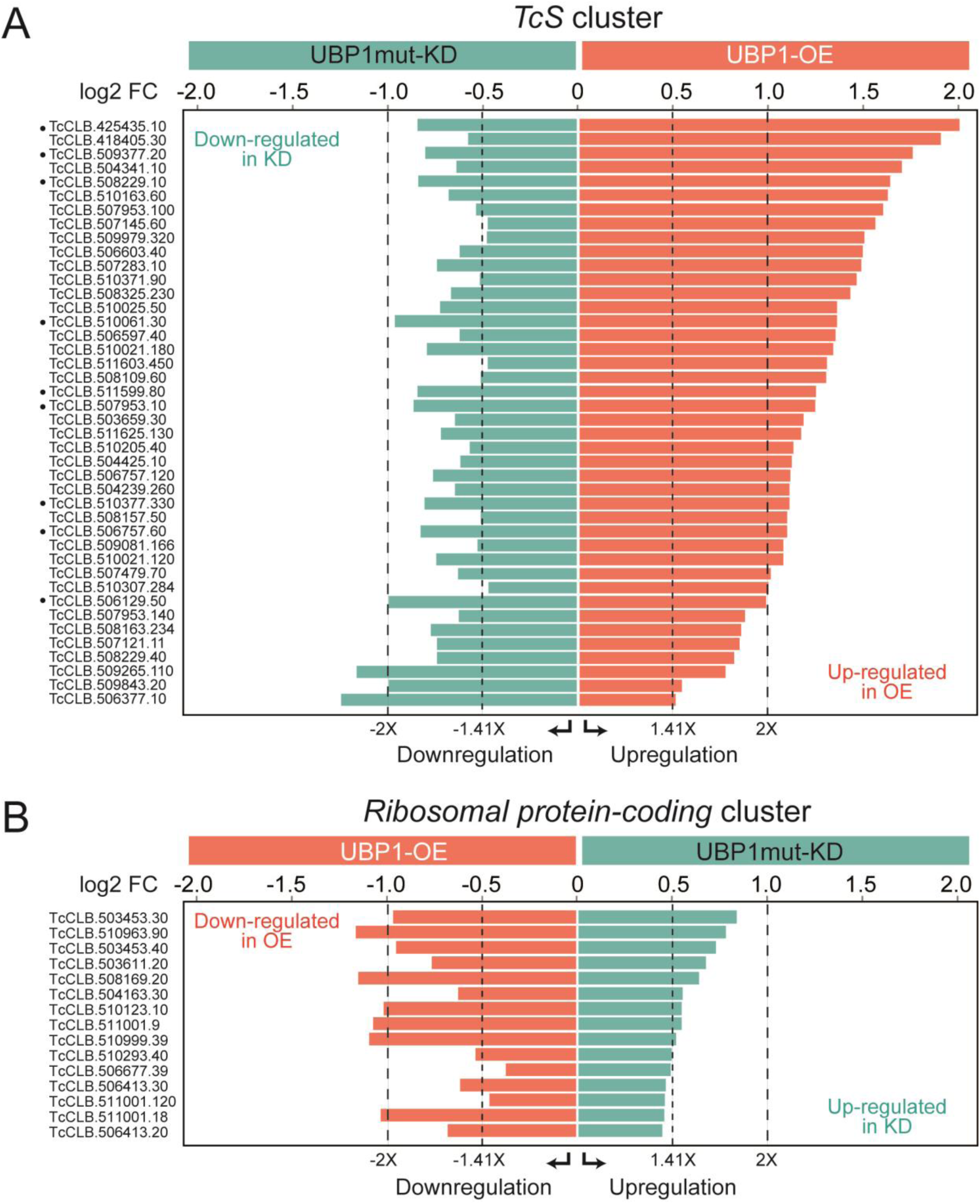
Bar charts displaying enrichment levels of transcripts encoding TcS and ribosomal proteins oppositely regulated by TcUBP1. Genes from tables 3 and 4 were identified as significant using an FDR-adjusted *p* value < 5% in both the OE and the KD samples with respect to the control WT population. *A*, *TcS* cluster (n= 42). *B*, *Ribosomal protein-coding* cluster (n = 15). The expression differences of ±1.41, and ±2 times are marked with different pointed lines.

**Table 3.** Stabilization of *trypomastigote glycoprotein-coding* genes mediated by TcUBP1.

| GeneID | Product name | log2<br>OE/WT | padj<br>OE | log2<br>KD/WT | padj<br>KD |
| --- | --- | --- | --- | --- | --- |
| TcCLB.508229.40 | trans-sialidase, Group V, putative | 0.82 | 1.5E-06 | -0.74 | 1.5E-04 |
| TcCLB.507953.140 | trans-sialidase, Group V, putative | 0.88 | 3.2E-11 | -0.62 | 7.3E-05 |
| TcCLB.425435.10 | trans-sialidase, putative (fragment) | 2.07 | 3.9E-18 | -0.84 | 1.0E-02 |
| TcCLB.508157.50 | trans-sialidase, putative | 1.12 | 4.1E-10 | -0.49 | 3.8E-02 |
| TcCLB.506129.50 | trans-sialidase, Group IV, putative | 1.00 | 2.6E-04 | -1.00 | 9.8E-04 |
| TcCLB.507121.11 | trans-sialidase, putative (fragment) | 0.85 | 7.6E-04 | -0.74 | 2.4E-02 |
| TcCLB.507953.100 | trans-sialidase, Group V, putative | 1.64 | 7.0E-22 | -0.52 | 1.8E-02 |
| TcCLB.510061.30 | trans-sialidase, Group V, putative | 1.39 | 2.7E-12 | -0.97 | 1.8E-04 |
| TcCLB.506377.10 | trans-sialidase, putative | 0.50 | 5.0E-03 | -1.27 | 1.2E-10 |
| TcCLB.508109.60 | trans-sialidase, Group V, putative | 1.33 | 2.6E-22 | -0.49 | 3.8E-03 |
| TcCLB.506603.40 | trans-sialidase, Group V, putative | 1.53 | 3.4E-25 | -0.61 | 7.8E-04 |
| TcCLB.508229.10 | trans-sialidase, putative | 1.69 | 2.0E-10 | -0.84 | 2.3E-02 |
| TcCLB.506757.60 | trans-sialidase, Group V, putative | 1.11 | 2.1E-21 | -0.83 | 2.2E-10 |
| TcCLB.508163.234 | trans-sialidase, putative | 0.86 | 5.1E-07 | -0.77 | 2.0E-04 |
| TcCLB.509081.166 | trans-sialidase, putative (fragment) | 1.09 | 7.7E-09 | -0.51 | 3.9E-02 |
| TcCLB.510025.50 | trans-sialidase, Group V, putative | 1.39 | 2.7E-24 | -0.72 | 7.7E-06 |
| TcCLB.504239.260 | trans-sialidase, Group V, putative | 1.13 | 2.6E-10 | -0.64 | 3.5E-03 |
| TcCLB.510377.330 | trans-sialidase, Group V, putative | 1.13 | 6.8E-18 | -0.81 | 9.6E-08 |
| TcCLB.510307.284<sup>#</sup> | trans-sialidase, Group IV, putative | 1.01 | 2.4E-06 | -0.45 | 9.8E-02 |
| TcCLB.504341.10 | trans-sialidase, Group IV, putative | 1.75 | 2.4E-14 | -0.63 | 3.7E-02 |
| TcCLB.507145.60 | trans-sialidase, Group V, putative | 1.60 | 5.6E-27 | -0.46 | 2.4E-02 |
| TcCLB.509843.20 | trans-sialidase, Group VII, putative | 0.53 | 4.8E-04 | -1.01 | 6.9E-10 |
| TcCLB.503659.30 | trans-sialidase, putative | 1.20 | 9.3E-12 | -0.64 | 4.5E-03 |
| TcCLB.511599.80 | trans-sialidase, Group V, putative | 1.28 | 4.4E-17 | -0.84 | 4.7E-06 |
| TcCLB.509377.20<sup>#</sup> | trans-sialidase, Group VII, putative | 1.81 | 5.5E-07 | -0.80 | 8.2E-02 |
| TcCLB.418405.30<sup>#</sup> | trans-sialidase, putative | 1.97 | 8.7E-22 | -0.56 | 5.3E-02 |
| TcCLB.509979.320 | trans-sialidase, Group V, putative | 1.54 | 5.8E-33 | -0.46 | 3.0E-03 |
| TcCLB.509265.110 | trans-sialidase, Group IV, putative | 0.77 | 2.5E-02 | -1.18 | 1.8E-03 |
| TcCLB.506757.120 | trans-sialidase, Group V, putative | 1.13 | 3.2E-21 | -0.76 | 1.1E-08 |
| TcCLB.507479.70 | trans-sialidase, Group IV, putative | 1.02 | 1.8E-05 | -0.62 | 4.0E-02 |
| TcCLB.506597.40 | trans-sialidase, Group V, putative | 1.38 | 3.3E-35 | -0.61 | 3.6E-06 |
| TcCLB.510205.40 | trans-sialidase, Group V, putative | 1.15 | 1.6E-14 | -0.56 | 2.1E-03 |
| TcCLB.511625.130 | trans-sialidase, Group V, putative | 1.19 | 6.2E-18 | -0.72 | 9.7E-06 |
| TcCLB.511603.450 | trans-sialidase, Group V, putative | 1.34 | 3.1E-15 | -0.46 | 4.4E-02 |
| TcCLB.510021.180 | trans-sialidase, Group V, putative | 1.37 | 3.4E-25 | -0.79 | 2.8E-07 |
| TcCLB.510371.90 | trans-sialidase, Group V, putative | 1.50 | 1.9E-29 | -0.50 | 2.6E-03 |
| TcCLB.504425.10 | trans-sialidase, Group III, putative | 1.14 | 1.1E-10 | -0.61 | 3.2E-03 |
| TcCLB.507953.10 | trans-sialidase, Group V, putative | 1.27 | 8.9E-09 | -0.87 | 2.6E-03 |
| TcCLB.510021.120 | trans-sialidase, Group V, putative | 1.09 | 2.8E-21 | -0.74 | 1.2E-08 |
| TcCLB.507283.10 | trans-sialidase, Group V, putative | 1.52 | 1.9E-41 | -0.74 | 2.9E-08 |
| TcCLB.510163.60 | trans-sialidase, Group V, putative | 1.67 | 1.8E-55 | -0.67 | 7.1E-08 |
| TcCLB.508325.230 | trans-sialidase, Group IV, putative | 1.46 | 9.7E-17 | -0.66 | 1.2E-03 |

42 significant UBPI-mRNA targets coding for *TcS* superfamily members oppositely regulated in OE (stabilized) and KD (destabilized) parasites using a combined criterion of log<sub>2</sub> fold change OE/WT > 0.45 and log<sub>2</sub> fold change KD/WT < -0.45 (1.37X, FDR-adjusted *p* value < 5%).
<sup>#</sup>Genes with FDR-adjusted *p* value in KD parasites < 10%.

**Table 4.**
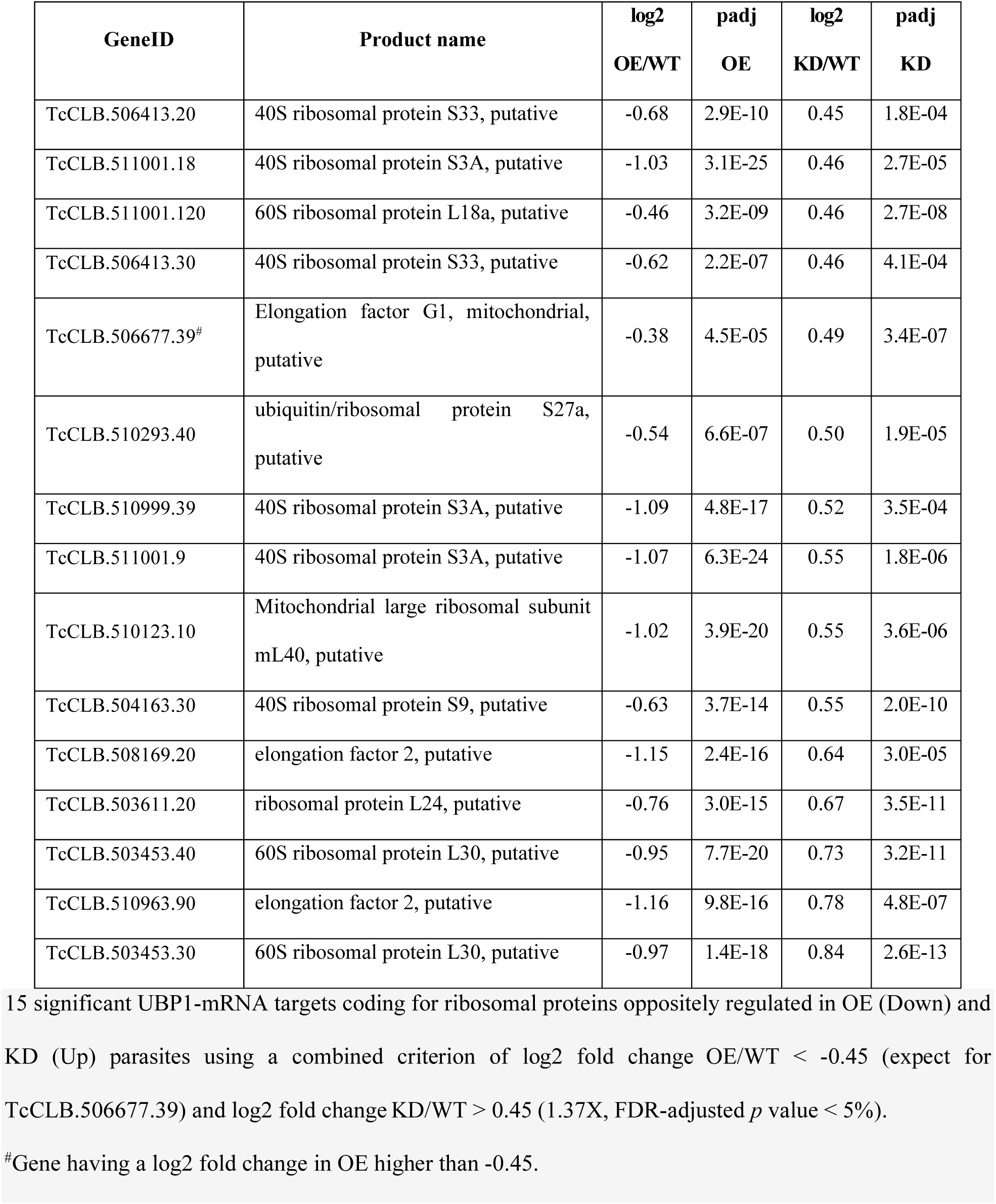
Destabilization of *ribosomal protein-coding* genes mediated by TcUBP1.

| GeneID | Product name | log2<br>OE/WT | padj<br>OE | log2<br>KD/WT | padj<br>KD |
| --- | --- | --- | --- | --- | --- |
| TcCLB.506413.20 | 40S ribosomal protein S33, putative | -0.68 | 2.9E-10 | 0.45 | 1.8E-04 |
| TcCLB.511001.18 | 40S ribosomal protein S3A, putative | -1.03 | 3.1E-25 | 0.46 | 2.7E-05 |
| TcCLB.511001.120 | 60S ribosomal protein L18a, putative | -0.46 | 3.2E-09 | 0.46 | 2.7E-08 |
| TcCLB.506413.30 | 40S ribosomal protein S33, putative | -0.62 | 2.2E-07 | 0.46 | 4.1E-04 |
| TcCLB.506677.39<sup>#</sup> | Elongation factor G1, mitochondrial,<br>putative | -0.38 | 4.5E-05 | 0.49 | 3.4E-07 |
| TcCLB.510293.40 | ubiquitin/ribosomal protein S27a,<br>putative | -0.54 | 6.6E-07 | 0.50 | 1.9E-05 |
| TcCLB.510999.39 | 40S ribosomal protein S3A, putative | -1.09 | 4.8E-17 | 0.52 | 3.5E-04 |
| TcCLB.511001.9 | 40S ribosomal protein S3A, putative | -1.07 | 6.3E-24 | 0.55 | 1.8E-06 |
| TcCLB.510123.10 | Mitochondrial large ribosomal subunit<br>mL40, putative | -1.02 | 3.9E-20 | 0.55 | 3.6E-06 |
| TcCLB.504163.30 | 40S ribosomal protein S9, putative | -0.63 | 3.7E-14 | 0.55 | 2.0E-10 |
| TcCLB.508169.20 | elongation factor 2, putative | -1.15 | 2.4E-16 | 0.64 | 3.0E-05 |
| TcCLB.503611.20 | ribosomal protein L24, putative | -0.76 | 3.0E-15 | 0.67 | 3.5E-11 |
| TcCLB.503453.40 | 60S ribosomal protein L30, putative | -0.95 | 7.7E-20 | 0.73 | 3.2E-11 |
| TcCLB.510963.90 | elongation factor 2, putative | -1.16 | 9.8E-16 | 0.78 | 4.8E-07 |
| TcCLB.503453.30 | 60S ribosomal protein L30, putative | -0.97 | 1.4E-18 | 0.84 | 2.6E-13 |

15 significant UBP1-mRNA targets coding for ribosomal proteins oppositely regulated in OE (Down) and KD (Up) parasites using a combined criterion of log2 fold change OE/WT < -0.45 (except for TcCLB.506677.39) and log2 fold change KD/WT > 0.45 (1.37X, FDR-adjusted *p* value < 5%).
<sup>#</sup>Gene having a log2 fold change in OE higher than -0.45.

By contrast, the main cluster among the downregulated genes previously reported in the OE parasite transcriptome was that coding for ribosomal proteins [16]. Within this functional category, we found 15 genes whose abundance varied in a coordinated way in both OE and KD conditions (|log2 fold change| >0.45, FDR-adjusted *p* value < 5%), **Fig 8B** and **Table 4**). These results confirm that the mRNA abundance of several genes coding for ribosomal proteins is inversely influenced by TcUBP1 levels, either decreasing when UBP1 protein levels are high or increasing when UBP1 protein levels are low. Overall, the analysis of the consistent mRNAs that were affected in both conditions in opposite ways revealed the genes whose regulation is directly governed by TcUBP1. These were mRNAs coding for cell-surface glycoproteins, augmented by UBP1; and transcripts coding for ribosomal and a group of hypothetical proteins, diminished by UBP1.

### The gene expression profile observed in UBP1-OE parasites resembles that of stationary-phase epimastigotes, while UBP1mut-KD exhibits a profile more similar to a replicative stage

In this section, we utilized the RNA-Seq data from UBP1mut-KD to compare its expression profile with those of the four stages of *T. cruzi* [5, 39]. We calculated the percentage of regulated transcripts in KD parasites among the most up- or downregulated genes in a pairwise comparison between the epimastigotes (Epi), amastigotes (Ama), metacyclic trypomastigotes (MT) and trypomastigotes (Trypo) stages (**Fig 9A**). It was found that the KD transcriptome showed greater similarity, albeit low, with the Ama/Trypo and Epi/Trypo datasets (genes overrepresented in Ama or Epi with respect to Trypo). This can be visualized by the different statistically significant colored clusters in the heatmap depicted in **Figure 9A**.

**Fig 9.**
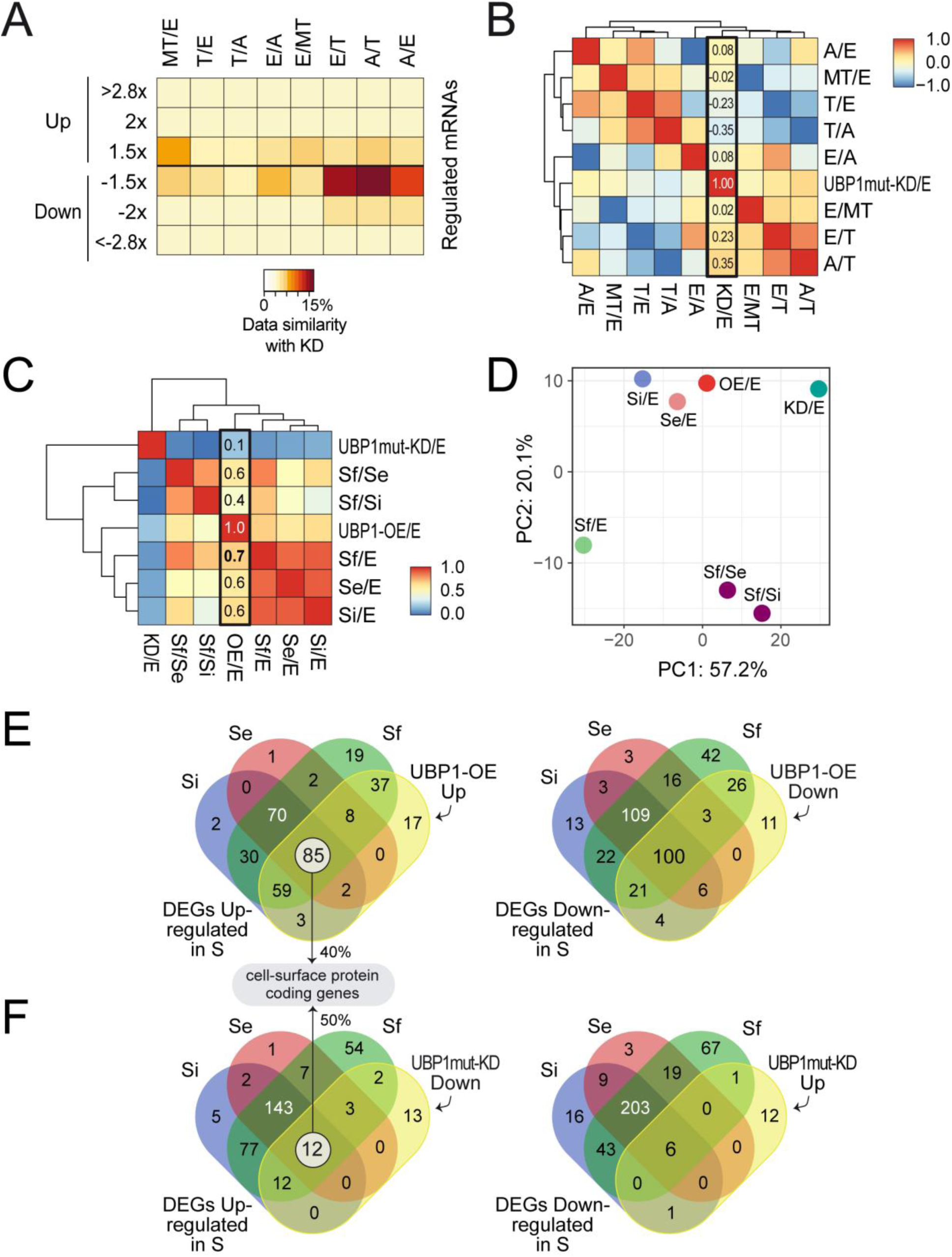
Comparison of UBP1-OE and UBP1mut-KD transcriptomes with distinct RNA-Seq datasets of *T. cruzi*. ***A***, heatmap representation of the percentages of shared genes between UBP1mut-KD and different pairwise comparisons. The brown/orange color indicates greater overlap, whereas the yellow color indicates less overlap. ***B***, heatmap representation of the Pearson correlation between KD samples with different *T. cruzi* stages. ***C***, heatmap representation of the Pearson correlation between OE and KD samples with different epimastigote stationary points: Se (early stationary, day 17 vs. 7), Si (intermediate stationary, day 21 vs. 7), Sf (final stationary, day 28 vs. 7), Sf vs. Si and Sf vs. Se. The red/orange color indicates a high correlation, whereas the blue/yellow color indicates a low correlation. ***D***, principal component (PC) analysis plot displaying the same samples as in *A*, along PC1 and PC2, which describe 57% and 20% of the variability, respectively. PC analysis was applied to 681 genes with log2 fold change data for all the pairwise comparisons. ***E***, Venn diagrams of Se, Si, Sf and UBP1-OE upregulated (left) and downregulated (right) genes. ***F***, Venn diagrams of DEGs upregulated in Se, Si and Sf with UBP1mut-KD downregulated genes (left) and *vice versa* (right).

Besides, we obtained fold change values for 2337 genes from the RNA-Seq experiments. These expression values were then used to calculate the Pearson correlation of all the samples, revealing a generally low correlation of the KD/Epi sample with the rest of the variables (column boxed in **Fig 9B**). Again, positive correlations were noted with Ama/Trypo (0.3471) and Epi/Trypo (0.2323), suggesting a weak but direct association. No significant correlation was found between KD and any of the remaining RNA-Seq experiments (**Fig 9B**). This analysis showed that the expression profile of the UBP1-KD population is more similar to that of the replicative stages than to that of the infective stages.

We next performed a comparative transcriptomic analysis using the recently published RNA-Seq data from Smircich *et al.* [40] to compare the expression profiles of TcUBP1-OE and KD parasites with those of *T. cruzi* epimastigotes in prolonged culture conditions (maintained until days 14, 21 and 28). For this, we obtained fold change values for 681 genes from the RNA-Seq experiments. The transcriptomic data groups in this analysis were: UBP1-OE, UBP1mut-KD, three successive growing selected points: early (Se, day 14), intermediate (Si, day 21) and final epimastigote stationary phase (Sf, day 28) relative to exponentially growing epimastigotes (E, day 7), and Sf compared to Se or Si points (Sf *versus* Se/Si). The correlation analysis showed that UBP1-OE parasites had the highest correlation with the datasets of epimastigotes in the stationary phase (r = 0.6640 for Sf, r = 0.6307 for Si, and r = 0.5735 for Se). No significant correlation was found between the UBP1-OE and KD RNA-Seq datasets, or between the KD sample and any of the other epimastigote stationary-phase datasets (**Fig 9C**).

The RNA-Seq expression table was also used to perform a principal component analysis (PCA) to compare the dispersion of the different datasets. The horizontal and vertical axes describe 77.3% of the variability, and, considering both PC1 and PC2, the UBP1-OE sample is distinctly located closer to the Se and Si experiments than to UBP-KD. Thus, this analysis showed that the expression profile of the UBP1-OE population is more similar to that of the stationary-phase epimastigotes than to that of the UBP1mut-KD condition (**Fig 9D**). The PCA and correlation between the different experiments analyzed showed that the expression profile of UBP1-overexpressing parasites resembles that of the epimastigote long-lasting growth points.

We focused on the analysis of shared DEGs in prolonged cultures of epimastigotes (Se∩Si∩Sf, hereinafter named as ST) with UBP1-OE or UBP1mut-KD parasites using a |log2 fold change| > 0.58 and FDR-adjusted *p* value < 0.10. For the UBP1-OE comparison, we filtered genes simultaneously up- or downregulated in both the ST and UBP1-OE groups (**Fig 9E**). For the UBP1mut-KD comparison, we focused on genes downregulated in KD and upregulated in ST (**Fig 9F**, left) or *vice versa* (**Fig 9F**, right). Regarding the ST∩OE DEGs, the upregulated list included 85 genes (**Fig 9E**, left and **S4 File**, *sheet A*), of which 40% code for trypomastigote cell surface-associated glycoproteins. The downregulated list contains 100 genes coding for proteins related to mitochondria or involved in translation and ribosome binding (**Fig 9E**, right and **S4 File**, *sheet B*). For ST∩KD, we found 12 genes within the DEGs downregulated in UBP1mut-KD and upregulated in ST, where six hits were related to surface glycoproteins of the trypomastigote form (**Fig 9F**, left and **S4 File**, *sheet C*). Lastly, six genes were obtained when intersecting the DEGs upregulated in UBP1mut-KD and downregulated in ST (**Fig 9F**, right and **S4 File**, *sheet D*). While these proteins have distinct functions, some may be related to common cellular processess such as stress response and protein degradation. Overall, the gene expression profile observed in *T. cruzi* UBP1-OE samples shared with the epimastigotes in starving conditions is compatible with a transitional parasite stage from replicative into infective metacyclic trypomastigotes in the insect host, while in the KD parasites, what is observed is that the transcriptomic signature more closely resembles that of replicative forms.

## DISCUSSION

Evolutionary conserved RBP networks and ncRNAs regulate the synthesis of populations of transcripts encoding proteins that participate in a common cellular process [reviewed in 41]. Trypanosomatid RBPs play a crucial role in orchestrating parasite differentiation, by coordinating the precise timing of developmental processes [11, 14, 15, 42, 43]. This holds for TcUBP1, a protein expressed across all developmental stages of the parasite. TcUBP1 functions by stabilizing or destabilizing myriad mRNAs, depending on the interaction with other stage-specific components of ribonucleoprotein complexes throughout the parasite life cycle [10]. Our previous findings revealed that overexpression of TcUBP1 in insect-dwelling epimastigotes enhanced the abundance of *TcS* transcripts and altered their subcellular localization to a perinuclear region [36, 44]. These observations were further supported by a shift toward mRNA expression characteristic of infective trypomastigotes [16, 36]. These findings agree with previous reports indicating a three-fold increase in TcUBP1 protein levels during parasite differentiation into metacyclic trypomastigotes [45].

In this study, we briefly present the methodology by which we obtained epimastigotes CL Brener UBP1mut-KD (**Fig 1**). Analysis of RNA-Seq data from KD samples confirmed that mapped reads for *TcUBP1* were interrupted at position 18, where the hygromycin CDS from the donor sequence was inserted, as outlined in the CRISPR-Cas9 experiment (**S2A Fig**). Western blotting allowed detecting a band of ∼24 kDa, with notably reduced expression. Worth noting, through cDNA analysis, we obtained a chimeric sequence with a 5′-end fragment from *UBP2* and a 3′-end segment from *UBP1*. Both open reading frames are encoded in the same polycistronic gene cluster, with the *UBP2* gene upstream of the *UBP1* gene [23], and given the diverse splicing processing that this molecule must undergo, it is feasible to hypothesize that a particular RNA rearrangement could have been carried out to potentially reverse the *TcUBP1* knockout. Given that the orthologous RRM-motif proteins TbUBP1 and TbUBP2 are essential for normal trypanosome growth in *T. brucei* [46], it is plausible to assume that this condition applies to *T. cruzi* as well.

The evidence shown in this paper confirms that TcUBP1 works by affecting the stability of its mRNA targets. TcUBP1 overexpression in the epimastigote cells led to a significant increase in the abundance of 793 genes and a downregulation of 371 genes [16]. In the population of UBP1mut-KD parasites, a significant decrease of 426 genes and an increase of 276 genes was observed (|log2 fold change| > 0.58, FRD-adjusted *p* value < 5%) (S2 File). Thus, the results of the present study reinforce the idea that the global balance of the TcUBP1 function on the epimastigote cells is mostly stabilization (**Fig 4**). By analyzing the transcriptome of these parasites using RNA-Seq experiments, we identified clusters having partly opposite expression patterns, increasing their expression in the UBP1-OE parasites and decreasing in the UBP1mut-KD population (**S3 File** and **S5 Fig**). Among the trypomastigote-stage specific genes, 16 were also upregulated in the transcriptome or translatome of metacyclic trypomastigotes [5] (see **S7A Fig**). Notably, 13 out of these 16 genes were trypomastigote-associated cell surface proteins (**S7B Fig**). Additionally, six genes were found at the intersection of all the analyzed datasets. Of these six, four are proteins belonging to the TcS superfamily (marked in red in **S7 Fig**).

Conversely, another cluster of genes related to translation showed increased expression in the UBP1mut-KD condition and decreased expression in the UBP1-OE condition. This change in abundances showed that numerous mRNAs differentially expressed in the transcriptome of UBP1mut-KD parasites are regulated in a contrasting manner with respect to those of the UBP1-OE population. The observed pattern of decreased expression in the OE sample and increased expression in the UBP1mut-KD sample suggests a potential involvement of these proteins in the replicative stages of the parasite (we identified six genes coding for hypothetical proteins in **Table 1**). In this context, through the use of a homemade bioinformatic algorithm, we successfully identified the TcCLB.503453.10 gene (HYPO_3) as the Complex I NDUFA5 subunit family (available at https://github.com/sradiouy/DARK). This suggests a heightened relevance during the replicative stages of the parasite, due to the role of this protein in energy production.

Our comparative transcriptome analysis revealed that the expression pattern of the UBP1-OE population closely resembles that of epimastigotes in the stationary phase, while it differs significantly from the UBP1mut-KD condition (**Fig 9**). These results highlight a substantial overlap between UBP1-OE and epimastigote stationary phase DEGs, supporting the notion that UBP1-overexpressing parasites exhibit characteristics of a transitional parasite form between epimastigotes and metacyclic trypomastigotes in the insect host. The transition from replicative to infective-type cell also requires a decrease in the rate of translation, and we currently know that this step is regulated by the expression and phosphorylation levels of the initiation factor eIF2α [20]. The knockout of TcK2 protein kinase, which phosphorylates *T. cruzi* eIF2α, showed lower release of cell-derived trypomastigotes than the control [47]. Moreover, the overexpression of eIF2α in amastigotes increases translation levels and decreases differentiation to infective trypomastigotes [20]. In the same line, overexpression of UBP1 in a replicative stage decreases the abundance of numerous transcripts coding for ribosomal proteins, probably withdrawing translation global levels and triggering an infective-type expression profile. So far, our results evidence once more the close relationship between regulatory posttranscriptional mechanisms and parasite differentiation, where TcUBP1 stands out as a central factor. In this regard, and in line with our results, it has been reported that mice immunized with parasites mutants for active trans-sialidases, which show less differentiation to cell-derived trypomastigotes, were fully protected against a challenge infection with the virulent *T. cruzi* Y strain [48]. Since these surface glycoproteins are part of a superfamily composed of numerous genes, studying the role of an upstream regulator, such as TcUBP1, in replicative amastigote cells is promising. Therefore, future research focusing on mutated TcUBP1 may pave the way for the development of new approaches to prevent the spread of infection.

## ACKNOWLEDGMENTS

We are indebted to Liliana Sferco and Agustina Chidichimo for parasite cultures. The pROCK-Cas9-GFP plasmid was gently provided by Dr. Santuza Teixeira.

## ABBREVIATIONS AND NOMENCLATURE

RBP: RNA-binding protein
TcUBP1: *T. cruzi* U-rich RBP 1
RRM: RNA-recognition motif
TcS: trans-sialidase/trans-sialidase-like
OE: overexpression
KD: knockdown
WT: wildtype
FDR: false discovery rate
GO: gene ontology
PCA: principal component analysis
Epi: epimastigote
MT: metacyclic trypomastigote
Ama: amastigote
Trypo: cell-derived trypomastigote

## Notes

### Competing Interest Statement

The authors have declared no competing interest.

## REFERENCES

1. Radío S, Fort RS, Garat B, Sotelo-Silveira J, Smircich P. UTRme: A Scoring-Based Tool to Annotate Untranslated Regions in Trypanosomatid Genomes. Front Genet. 2018;9: 671.

2. Smircich P, Forteza D, El-Sayed NM, Garat B. Genomic analysis of sequence-dependent DNA curvature in Leishmania. PLoS One. 2013;8: e63068.

3. Callejas-Hernández F, Gutierrez-Nogues Á, Rastrojo A, Gironès N, Fresno M. Analysis of mRNA processing at whole transcriptome level, transcriptomic profile and genome sequence refinement of Trypanosoma cruzi. Sci Rep. 2019;9: 17376.

4. Barbosa RL, da Cunha JPC, Menezes AT, Melo R de FP, Elias MC, Silber AM, et al. Proteomic analysis of Trypanosoma cruzi spliceosome complex. J Proteomics. 2020;223: 103822.

5. Smircich P, Eastman G, Bispo S, Duhagon MA, Guerra-Slompo EP, Garat B, et al. Ribosome profiling reveals translation control as a key mechanism generating differential gene expression in Trypanosoma cruzi. BMC Genomics. 2015;16: 443.

6. Jensen BC, Ramasamy G, Vasconcelos EJR, Ingolia NT, Myler PJ, Parsons M. Extensive stage-regulation of translation revealed by ribosome profiling of Trypanosoma brucei. BMC Genomics. 2014;15: 911.

7. Vasquez J-J, Hon C-C, Vanselow JT, Schlosser A, Siegel TN. Comparative ribosome profiling reveals extensive translational complexity in different Trypanosoma brucei life cycle stages. Nucleic Acids Res. 2014;42: 3623–3637.

8. Pastro L, Smircich P, Di Paolo A, Becco L, Duhagon MA, Sotelo-Silveira J, et al. Nuclear Compartmentalization Contributes to Stage-Specific Gene Expression Control in. Front Cell Dev Biol. 2017;5: 8.

9. Fadda A, Ryten M, Droll D, Rojas F, Färber V, Haanstra JR, et al. Transcriptome-wide analysis of trypanosome mRNA decay reveals complex degradation kinetics and suggests a role for co-transcriptional degradation in determining mRNA levels. Mol Microbiol. 2014;94: 307–326.

10. De Gaudenzi JG, Noé G, Campo VA, Frasch AC, Cassola A. Gene expression regulation in trypanosomatids. Essays Biochem. 2011;51: 31–46.

11. Kolev NG, Ramey-Butler K, Cross GAM, Ullu E, Tschudi C. Developmental progression to infectivity in Trypanosoma brucei triggered by an RNA-binding protein. Science. 2012;338: 1352– 1353.

12. Jha BA, Gazestani VH, Yip CW, Salavati R. The DRBD13 RNA binding protein is involved in the insect-stage differentiation process of Trypanosoma brucei. FEBS Lett. 2015;589: 1966–1974.

13. Shi H, Butler K, Tschudi C. Differential expression analysis of transcriptome data of RBP6 induction in procyclics leading to infectious metacyclics and bloodstream forms. Data Brief. 2018;20: 978–980.

14. Alcantara MV, Kessler RL, Gonçalves REG, Marliére NP, Guarneri AA, Picchi GFA, et al. Knockout of the CCCH zinc finger protein TcZC3H31 blocks Trypanosoma cruzi differentiation into the infective metacyclic form. Mol Biochem Parasitol. 2018;221: 1–9.

15. Tavares TS, Mügge FLB, Grazielle-Silva V, Valente BM, Goes WM, Oliveira AER, et al. A zinc finger protein that is implicated in the control of epimastigote-specific gene expression and metacyclogenesis. Parasitology. 2021;148: 1171–1185.

16. Sabalette KB, Sotelo-Silveira JR, Smircich P, De Gaudenzi JG. RNA-Seq reveals that overexpression of TcUBP1 switches the gene expression pattern toward that of the infective form of Trypanosoma cruzi. J Biol Chem. 2023;299: 104623.

17. De Souza W, Barrias ES. May the epimastigote form of Trypanosoma cruzi be infective? Acta Trop. 2020;212: 105688.

18. Brener Z. Life cycle of Trypanosoma cruzi. Rev Inst Med Trop Sao Paulo. 1971;13: 171–178.

19. Radío S, Garat B, Sotelo-Silveira J, Smircich P. Upstream ORFs Influence Translation Efficiency in the Parasite. Front Genet. 2020;11: 166.

20. Castro Machado F, Bittencourt-Cunha P, Malvezzi AM, Arico M, Radio S, Smircich P, et al. EIF2α phosphorylation is regulated in intracellular amastigotes for the generation of infective Trypanosoma cruzi trypomastigote forms. Cell Microbiol. 2020;22: e13243.

21. De Gaudenzi J, Frasch AC, Clayton C. RNA-binding domain proteins in Kinetoplastids: a comparative analysis. Eukaryot Cell. 2005;4: 2106–2114.

22. De Gaudenzi JG, D’Orso I, Frasch ACC. RNA recognition motif-type RNA-binding proteins in Trypanosoma cruzi form a family involved in the interaction with specific transcripts in vivo. J Biol Chem. 2003;278: 18884–18894.

23. De Gaudenzi JG, Jäger AV, Izcovich R, Campo VA. Insights into the Regulation of mRNA Processing of Polycistronic Transcripts Mediated by DRBD4/PTB2, a Trypanosome Homolog of the Polypyrimidine Tract-Binding Protein. J Eukaryot Microbiol. 2016;63: 440–452.

24. D’Orso I, Frasch ACC. TcUBP-1, an mRNA Destabilizing Factor from Trypanosomes, Homodimerizes and Interacts with Novel AU-rich Element- and Poly(A)-binding Proteins Forming a Ribonucleoprotein Complex. Journal of Biological Chemistry. 2002. pp. 50520–50528. doi:10.1074/jbc.m209092200

25. Wippel HH, Inoue AH, Vidal NM, Costa JF da, Marcon BH, Romagnoli BAA, et al. Assessing the partners of the RBP9-mRNP complex in Trypanosoma cruzi using shotgun proteomics and RNA-seq. RNA Biol. 2018;15: 1106–1118.

26. Wippel HH, Malgarin JS, Inoue AH, da Veiga Leprevost F, Carvalho PC, Goldenberg S, et al. Unveiling the partners of the DRBD2-mRNP complex, an RBP in Trypanosoma cruzi and ortholog to the yeast SR-protein Gbp2. BMC Microbiology. 2019. doi:10.1186/s12866-019-1505-8

27. Noé G, De Gaudenzi JG, Frasch AC. Functionally related transcripts have common RNA motifs for specific RNA-binding proteins in trypanosomes. BMC Mol Biol. 2008;9: 107.

28. Li Z-H, De Gaudenzi JG, Alvarez VE, Mendiondo N, Wang H, Kissinger JC, et al. A 43-nucleotide U-rich element in 3’-untranslated region of large number of Trypanosoma cruzi transcripts is important for mRNA abundance in intracellular amastigotes. J Biol Chem. 2012;287: 19058–19069.

29. De Gaudenzi JG, Carmona SJ, Agüero F, Frasch AC. Genome-wide analysis of 3’-untranslated regions supports the existence of post-transcriptional regulons controlling gene expression in trypanosomes. PeerJ. 2013;1: e118.

30. Keene JD. RNA regulons: coordination of post-transcriptional events. Nat Rev Genet. 2007;8: 533– 543.

31. Keene JD, Lager PJ. Post-transcriptional operons and regulons co-ordinating gene expression. Chromosome Res. 2005;13: 327–337.

32. Bisogno LS, Keene JD. RNA regulons in cancer and inflammation. Curr Opin Genet Dev. 2018;48: 97–103.

33. Lander N, Bernal C, Diez N, Añez N, Docampo R, Ramírez JL. Localization and developmental regulation of a dispersed gene family 1 protein in Trypanosoma cruzi. Infect Immun. 2010;78: 231– 240.

34. Lander N, Chiurillo MA, Docampo R. Signaling pathways involved in environmental sensing in Trypanosoma cruzi. Mol Microbiol. 2021;115: 819–828.

35. Romagnoli BAA, Holetz FB, Alves LR, Goldenberg S. RNA Binding Proteins and Gene Expression Regulation in Trypanosoma cruzi. Frontiers in Cellular and Infection Microbiology. 2020. doi:10.3389/fcimb.2020.00056

36. Sabalette KB, Romaniuk MA, Noé G, Cassola A, Campo VA, De Gaudenzi JG. The RNA-binding protein TcUBP1 up-regulates an RNA regulon for a cell surface-associated glycoprotein and promotes parasite infectivity. J Biol Chem. 2019;294: 10349–10364.

37. Lee RTH, Ng ASM, Ingham PW. Ribozyme Mediated gRNA Generation for In Vitro and In Vivo CRISPR/Cas9 Mutagenesis. PLoS One. 2016;11: e0166020.

38. Cassola A, De Gaudenzi JG, Frasch AC. Recruitment of mRNAs to cytoplasmic ribonucleoprotein granules in trypanosomes. Mol Microbiol. 2007;65: 655–670.

39. Li Y, Shah-Simpson S, Okrah K, Belew A, Choi J, Caradonna K et al. Transcriptome remodeling in *Trypanosoma cruzi* and human cells during intracellular infection. Plos Pathog. 2016; 12:e100511.

40. Smircich P, Pérez-Díaz L, Hernández F, Duhagon MA, Garat B. Transcriptomic analysis of the adaptation to prolonged starvation of the insect-dwelling epimastigotes. Front Cell Infect Microbiol. 2023;13: 1138456.

41. Ho JJD, Man JHS, Schatz JH, Marsden PA. Translational remodeling by RNA-binding proteins and noncoding RNAs. Wiley Interdiscip Rev RNA. 2021;12: e1647.

42. Pérez-Díaz L, Correa A, Moretão MP, Goldenberg S, Dallagiovanna B, Garat B. The overexpression of the trypanosomatid-exclusive TcRBP19 RNA-binding protein affects cellular infection by Trypanosoma cruzi. Mem Inst Oswaldo Cruz. 2012;107: 1076–1079.

43. Romaniuk MA, Frasch AC, Cassola A. Translational repression by an RNA-binding protein promotes differentiation to infective forms in Trypanosoma cruzi. PLoS Pathog. 2018;14: e1007059.

44. Oliveira C, Holetz FB, Alves LR, Ávila AR. Modulation of Virulence Factors during Differentiation. Pathogens. 2022;12. doi:10.3390/pathogens12010032

45. de Godoy LMF, Marchini FK, Pavoni DP, Rampazzo R de CP, Probst CM, Goldenberg S, et al. Quantitative proteomics of Trypanosoma cruzi during metacyclogenesis. Proteomics. 2012;12: 2694– 2703.

46. Hartmann C, Benz C, Brems S, Ellis L, Luu V-D, Stewart M, et al. Small trypanosome RNA-binding proteins TbUBP1 and TbUBP2 influence expression of F-box protein mRNAs in bloodstream trypanosomes. Eukaryot Cell. 2007;6: 1964–1978.

47. Marcelino T de P, Fala AM, da Silva MM, Souza-Melo N, Malvezzi AM, Klippel AH, et al. Identification of inhibitors for the transmembrane Trypanosoma cruzi eIF2α kinase relevant for parasite proliferation. J Biol Chem. 2023;299: 104857.

48. Burle-Caldas G de A, Dos Santos NSA, de Castro JT, Mugge FLB, Grazielle-Silva V, Oliveira AER, et al. Disruption of Active Trans-Sialidase Genes Impairs Egress from Mammalian Host Cells and Generates Highly Attenuated Trypanosoma cruzi Parasites. MBio. 2022;13: e0347821.

