## Supporting information for "Transcriptomic analysis of N-terminal mutated *Trypanosoma cruzi* UBP1 knockdown underlines the importance of this RNA-binding protein in parasite development"

Running title: *Transcriptome of mutated T. cruzi UBP1-knockdown parasites*

*

**Keywords:** trypanosome; RNA-protein interaction; RNA-binding protein; gene regulation; RNA regulon.

---

Material included:

• S1-S4 Files

• S1-S7 Figures

• S1 Table

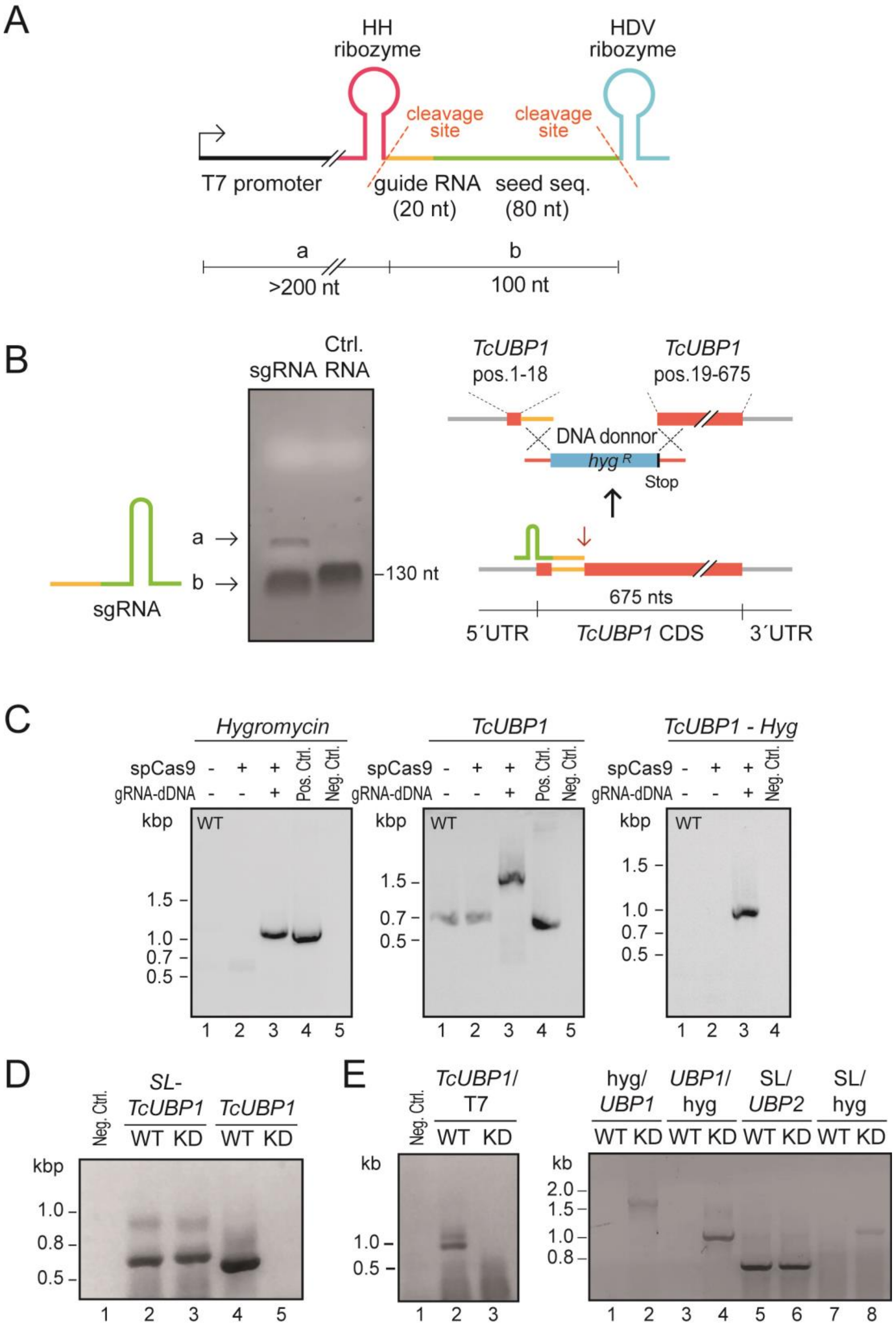
A
HH ribozyme
HDV ribozyme
cleavage site
cleavage site
T7 promoter
guide RNA (20 nt)
seed seq. (80 nt)
a
>200 nt
b
100 nt
B
sgRNA
Ctrl. RNA
a →
b →
130 nt
sgRNA
TcUBP1 pos.1-18
TcUBP1 pos.19-675
DNA donnor
hyg R
Stop
675 nts
5´UTR
TcUBP1 CDS
3´UTR
C
Hygromycin
TcUBP1
TcUBP1 - Hyg
spCas9
gRNA-dDNA
Pos. Ctrl.
Neg. Ctrl.
kbp
WT
1.5
1.0
0.7
0.5
1 2 3 4 5
1 2 3 4
D
Neg. Ctrl.
SL-TcUBP1
TcUBP1
WT KD WT KD
kbp
1.0
0.8
0.5
1 2 3 4 5
E
Neg. Ctrl.
TcUBP1/T7
WT KD
kb
1.0
0.5
1 2 3
hyg/UBP1
UBP1/hyg
SL/UBP2
SL/hyg
WT KD WT KD WT KD WT KD
kb
2.0
1.5
1.0
0.8
1 2 3 4 5 6 7 8

**S1 Fig. CRISPR-Cas9 genome editing of *TcUBP1*, PCR verification.** *A*, Scheme of the gRNA construct cloned into pEASY-T1 used for in vitro transcription. The T7 promoter region (black), HH ribozyme (magenta), the gRNA target complementary sequence (orange), the gRNA target sequence (green), and the cleavage sites of the ribosomes (dotted lines) are marked. *B*, Ethidium bromide-stained 2% agarose gel with denatured samples of the in vitro transcription gRNA product and a known-length RNA marker used as size control (Ctrl. RNA, 130 nt). The band of greater size corresponds to the gRNA upstream sequence (a), and one smaller band that is the gRNA sequence (b) are indicated with arrows; the sequence downstream to the gRNA is not observed in the gel due to its low size *(left)*, Scheme of the construct obtained after transfection with the complete system. The *hygromycin* resistance CDS was used as donor DNA (cyan) flanked by homologous sequences on both sides to the spCas9 site (tomato). The spCas9 cutting site is indicated within the coding sequence of *TcUBP1* (red arrow), *TcUBP1* CDS (tomato), 5’ and 3′-UTR of *TcUBP1*, and target sequence recognized by the gRNA (orange) (*right*). *C*, Agarose gel electrophoresis of PCR products from genomic DNA extracted from wildtype (WT) populations (lane 1), spCas9-GFP (lane 2) or populations transfected with the complete system (spCas9-gRNA-DNA donor) (lane 3) generated using forward and reverse-specific primers for *hygromycin*, for *TcUBP1*, or a combination of specific forward oligo for *TcUBP1* and reverse oligo for *hygromycin*. *D*, PCR products from cDNA of KD or WT parasites generated using a forward primer specific to the *Spliced leader* sequence and a reverse primer specific to the *TcUBP1* CDS; or with both specific primers for *TcUBP1* CDS. *E*, PCR products from cDNA of KD or WT parasites generated using a combination of forward/reverse primers specific for *TcUBP1*, *TcUBP2*, *SL*, *T7* or *hygromycin*. Molecular mass protein standards or DNA markers are indicated on the left. Pos. Ctrl., positive PCR control; Neg. Ctrl., negative PCR control. *SL*, spliced-leader sequence.

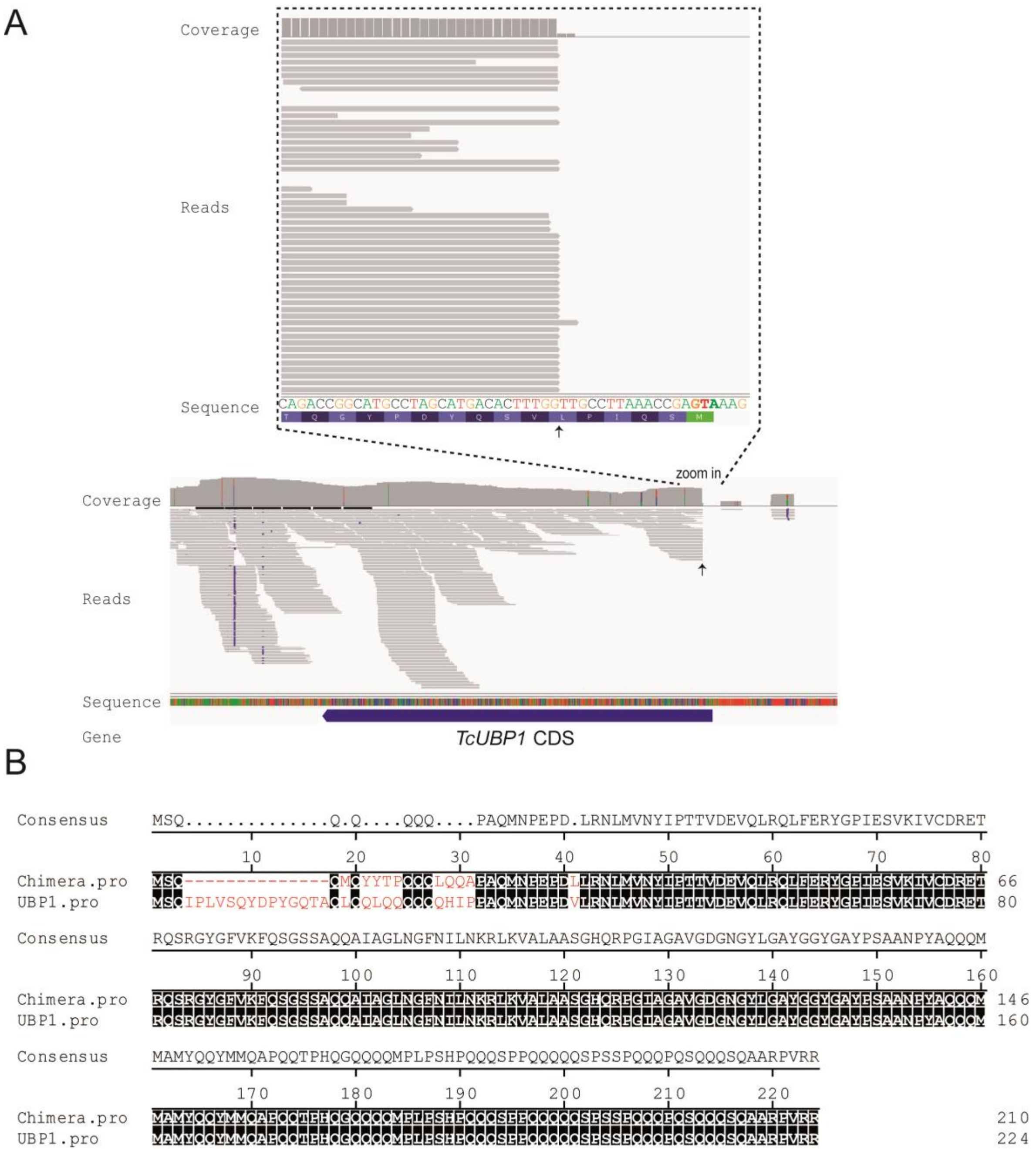


**S2 Fig. TcUBP1: visualization of RNA-seq reads and protein sequence alignment**. *A*, Mapped reads were visualized using IGV software. Coverage of reads is shown on the upper part of the IGV image. The scheme shows the 5′-end of *TcUBP1CDS* and the arrow indicates the position where the *hygromycin* gene was inserted. *B*, Pairwise sequence comparison between TcUBP1 and TcUBP1mut proteins obtained with the Jotun Hein Method. Sequences were aligned using Lasergene package (DNASTAR Inc.).

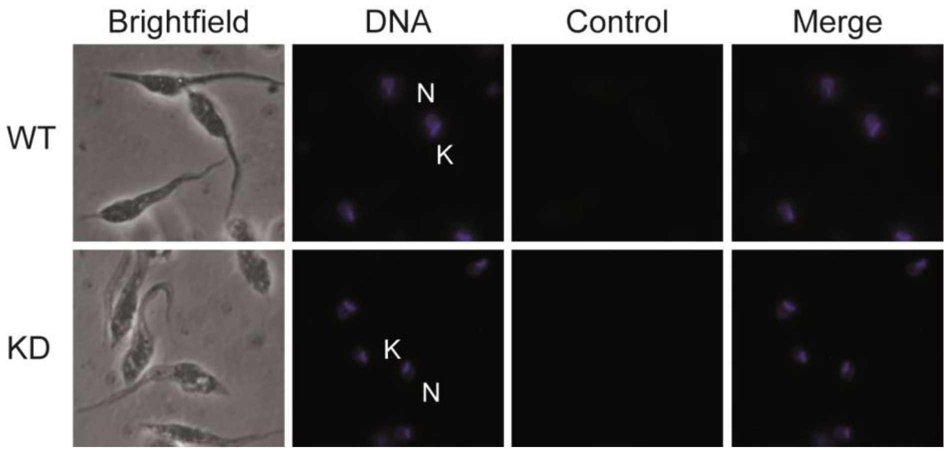


**S3 Fig. Microscopy images of control samples processed without the primary antibody.** Immunohistochemical staining of WT and KD samples processed without the primary antibody (red channel, control). In the DAPI panel (blue channel, DNA), the nuclear (N) and kinetoplast (K) DNA of *T. cruzi* are indicated.

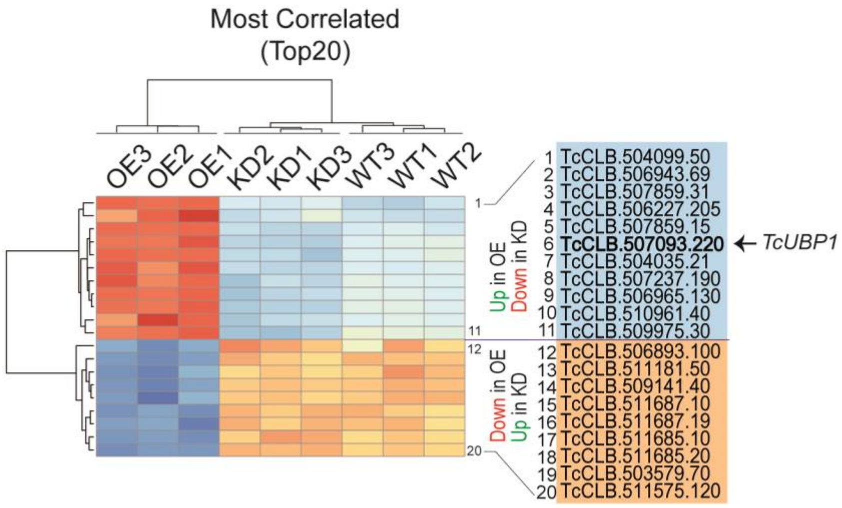


**S4 Fig. Most correlated samples (n = 20 genes).** The key is as for Fig 4*B*. The Z-score scale bar represents relative expression +/- SD from the mean.

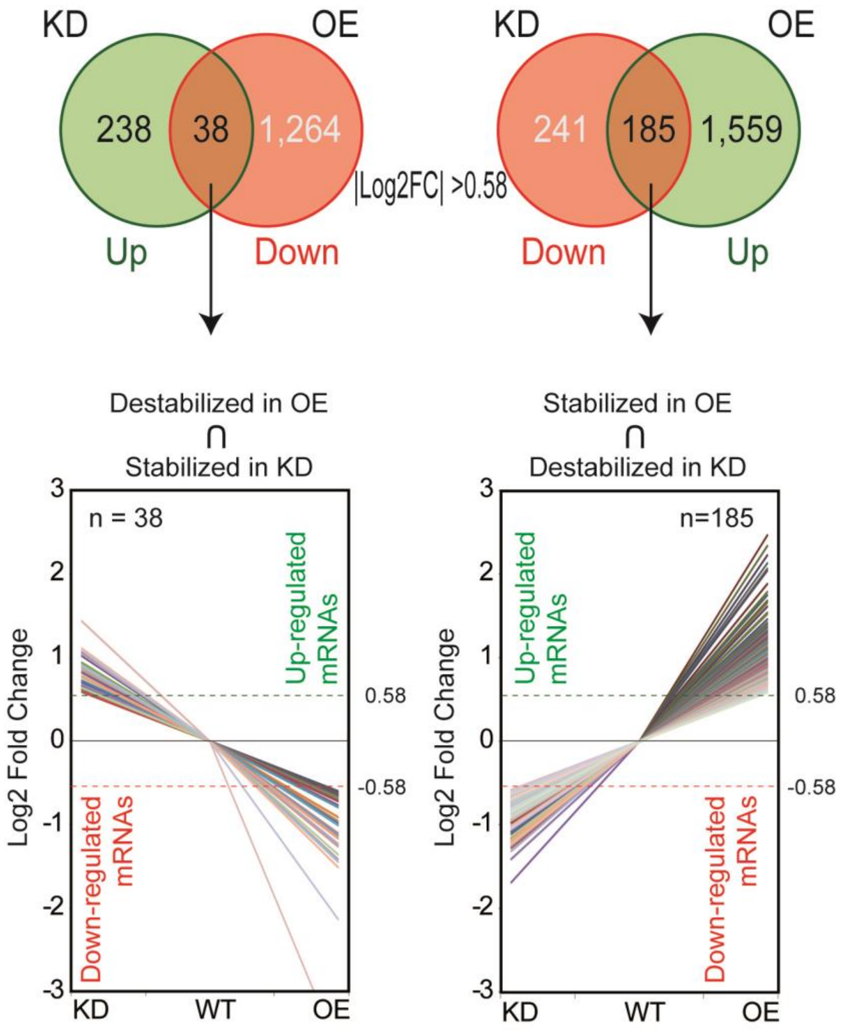


**S5 Fig. Transcripts whose abundance inversely responds to TcUBP1 expression levels.** The key is as for Fig. 5 but showing the number of genes 1.5-fold regulated in each condition OE and KD with respect to the WT control (|log2 fold change| >0.58). *TcUBP1* (TcCLB.507093.220) is not plotted, due to its extreme scale.

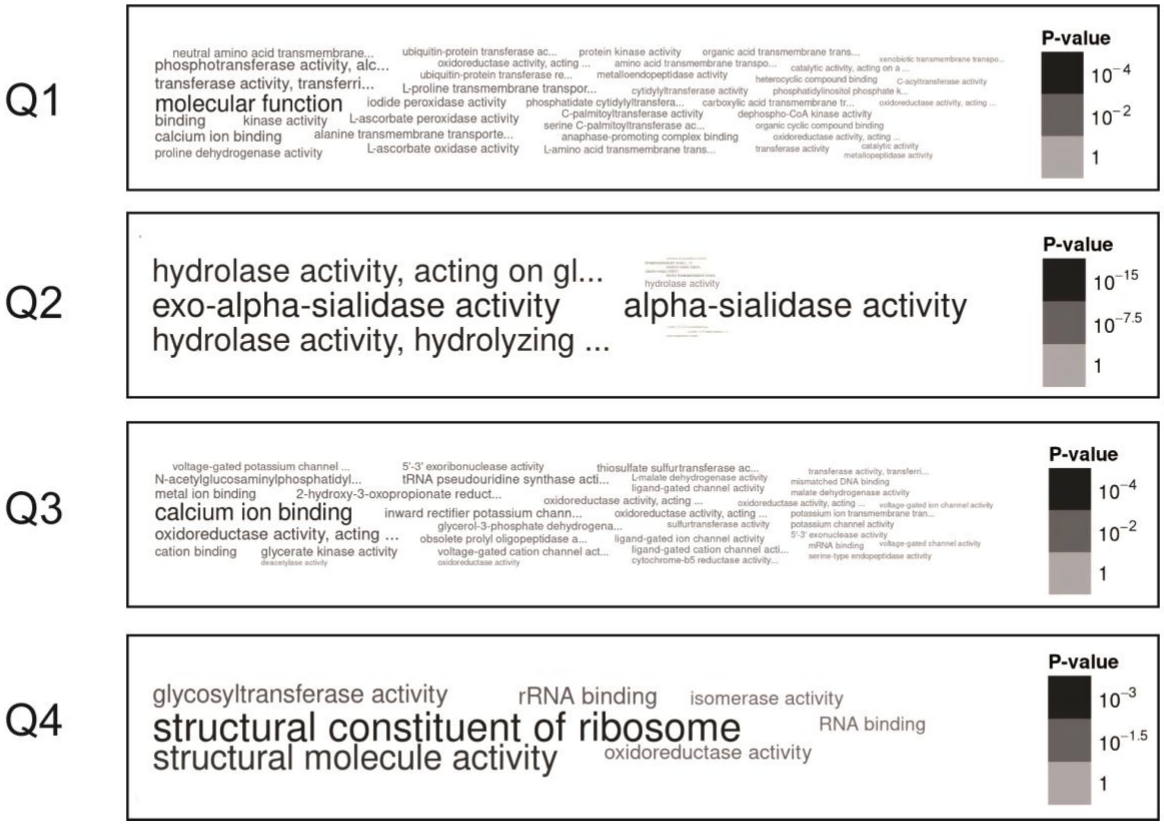


**S6 Fig. GO enrichment analysis of all the genes regulated in UBP1-OE *versus* KD parasites**. GO classification of DEGs using a criterion of at least 1.5-fold change (|log2 fold change| > 0.58); the graphs show a word cloud of terms for each quadrant of Figure 4*D* (only for GO domain: molecular function). Q1, n = 61 genes; Q2, n = 185 genes; Q3, n = 45 genes; Q4, n = 38 genes.

A

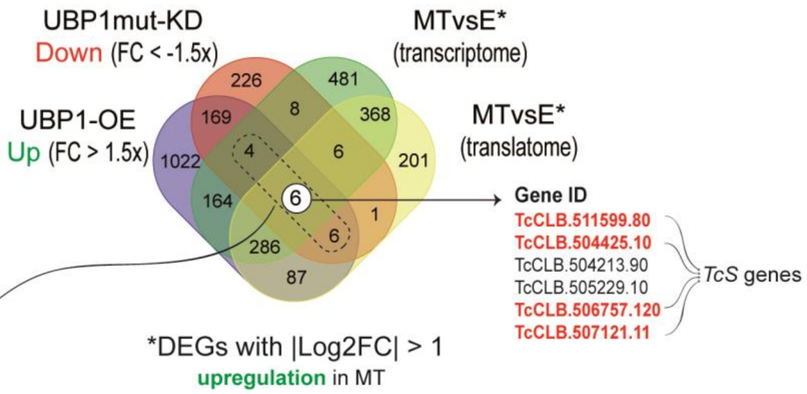


B

| Gene ID | Product Description |
|---|---|
| TcCLB.511675.3 | Trypomastigote, Alanine, Serine and Valine rich protein (TASV), subfamily C |
| TcCLB.504341.10 | trans-sialidase, Group IV, putative |
| **TcCLB.504425.10** | trans-sialidase, Group III, putative |
| TcCLB.505229.10 | Isoprenylcysteine alpha-carbonyl methylesterase, putative |
| TcCLB.506599.50 | Mucin-associated surface protein (MASP), subgroup S001 |
| TcCLB.506603.40 | trans-sialidase, Group V, putative |
| **TcCLB.507121.11** | trans-sialidase, putative (fragment) |
| **TcCLB.506757.120** | trans-sialidase, Group V, putative |
| TcCLB.504127.20 | hypothetical protein |
| TcCLB.504213.90 | inositol polyphosphate kinase-like protein, putative |
| TcCLB.507283.10 | trans-sialidase, Group V, putative |
| TcCLB.510025.50 | trans-sialidase, Group V, putative |
| **TcCLB.511599.80** | trans-sialidase, Group V, putative |
| TcCLB.507957.150 | Mucin-associated surface protein (MASP), subgroup S074 |
| TcCLB.508165.300 | Mucin-associated surface protein (MASP), subgroup S049 |
| TcCLB.510377.134 | Mucin-associated surface protein (MASP), subgroup S001 |

**S7 Fig. Venn diagram representing the total number of DEGs among UBP1-OE, UBP1mut-KD, metacyclic transcriptome and translatome.** *A*, UBP1-OE upregulated (log2 fold change >0.58), UBP1-mutKD downregulated (log2 fold change <-0.58), and transcriptome or translatome upregulated in MT (log2 fold change >1) are shown. *B*, list of the intersection of genes belonging to UBP1-OE (upregulated) ∩ UBP1mut-KD (downregulated) ∩ [MTvsE-transcriptome (upregulated) ∪ MTvsE-translatome (upregulated)]. In red, glycoprotein members of the *TcS* superfamily.

### S1 Table. Oligonucleotides used in this work.

| Gene | Primer name | Sequence (5′ to 3′) |
|---|---|---|
| *Hygromycin* | HYG-Fw | caaagaaaaTAGTTCAAACGAATTATGAAAAAGCCTGAACTCACC |
| | HYG-Rev | gagctcATTCCTTTGCCCTCGGACGA |
| *TcUBP1* | NH2-Fw | cgggatccATGAGCCAAATTGTTGTTTC |
| | NH2/AS | ATCGGGCTCGGGGTTCATCTG |
| | COOH-Rev | cgggatccTTACCTTCGAACAGGACGGGC |
| | TcSL | CTATTATTGATACAGTTTC |
| | Oligo T7 | TAATACGACTCACTATAGGG |
| | T7(dT) | TAATACGACTCACTATAGGG$(T)_{18}$ |
| *TcUBP2* | COOH-Rev | cggaattcCTACTGACGCGGCACCGACGG |
| sgRNA | 1. tRNA-C9xP-R-SacI | gagctcCTAGGAAAAAAAAAAAAAGGCCAC |
| | 2. UBP1(13)-gRNA1-Fw | gaggacgaaacgagtaagctcgtcGGATCGTACTGTGAAACCAAgttttagagctagaaatagcaagt |
| | 3. UBP1(13)-sgRNA1-Rev | gacgagcttactcgtttcgtcctcacggactcatcagGGATCGgctagcagggagagtgctaatcttc |
| | 4. HDV-C9xP-R-SpeI | AAAAAAAAAAAactagtCCCATTCGCCATG |
| Donor DNA | UBP1(13)-HYG-Fw | GTTTGAATTGTGGGAAATGAGCCAAATTCCGTTGatgaaaaagcctgaa |
| | UBP1(13)-HYG-Rev | GGGCAGTCTGGCCGTACGGATCGTACTGTGAAACctattcctttgccct |
| Sequence guide | Seq guide | GGATCGTACTGTGAAACCAACGG |

Restriction sites are underlined.

### SUPPLEMENTAL FILES

**S1 File. DEGs (>2X) up- or downregulated in UBP1mut-KD parasites.** List of up- and downregulated genes after UBP1 knockdown based on |log2 fold change| > 1 & FDR-adjusted *p* value < 0.05.

**S2 File. DEGs (>1.5X) up- or downregulated in UBP1mut-KD parasites.** List of up- and downregulated genes after UBP1 knockdown based on |log2 fold change| > 0.58 & FDR-adjusted *p* value < 0.05.

**Supplemental File S3.** Consistent genes affected in the opposite way in both OE and KD conditions (>1.5-fold or |Log2 fold change| = 0.58, FDR-adjusted *p* value < 0.05). Sheet *A*, 185 genes stabilized in OE parasites and destabilized in KD parasites; Sheet *B*, 38 genes stabilized in KD parasites and destabilized in OE parasites.

**Supplemental File S4.** Shared DEGs of the prolonged cultures of epimastigotes with UBP1-OE or UBP1mut-KD parasites (|log2 fold change| > 0.58 and FDR-adjusted p-value < 0.10). Sheet *A* (Table 1), 85 DEGs upregulated in OE and upregulated in Se∩Si∩Sf; Sheet *B* (Table 2), 100 DEGs downregulated in OE and downregulated in Se∩Si∩Sf; Sheet *C* (Table 3), 12 DEGs downregulated in UBP1mut-KD and upregulated in Se∩Si∩Sf; Sheet *D* (Table 4), 6 DEGs upregulated in UBP1mut-KD and downregulated in Se∩Si∩Sf.
